# Quantitative analysis of tumour spheroid structure

**DOI:** 10.1101/2021.08.05.455334

**Authors:** Alexander P Browning, Jesse A Sharp, Ryan J Murphy, Gency Gunasingh, Brodie Lawson, Kevin Burrage, Nikolas K Haass, Matthew J Simpson

## Abstract

Tumour spheroids are common *in vitro* experimental models of avascular tumour growth. Compared with traditional two-dimensional culture, tumour spheroids more closely mimic the avascular tumour microenvironment where spatial differences in nutrient availability strongly influence growth. We show that spheroids initiated using significantly different numbers of cells grow to similar limiting sizes, suggesting that avascular tumours have a limiting structure; in agreement with untested predictions of classical mathematical models of tumour spheroids. We develop a novel mathematical and statistical framework to study the structure of tumour spheroids seeded from cells transduced with fluorescent cell cycle indicators, enabling us to discriminate between arrested and cycling cells and identify an arrested region. Our analysis shows that transient spheroid structure is independent of initial spheroid size, and the limiting structure can be independent of seeding density. Standard experimental protocols compare spheroid size as a function of time; however, our analysis suggests that comparing spheroid structure as a function of overall size produces results that are relatively insensitive to variability in spheroid size. Our experimental observations are made using two melanoma cell lines, but our modelling framework applies across a wide range of spheroid culture conditions and cell lines.

## 1 Introduction

Three-dimensional tumour spheroids provide an accessible and biologically realistic *in vitro* model of early avascular tumour growth [1, 2]. Spheroids play a vital role in cancer therapy development, where the effect of a putative drug on spheroid growth is an indicator of efficacy [3–10]. In this context, reproducibility and uniformity in spheroid sizes is paramount [11–13], yet variability in the initial and final spheroid size is rarely accounted for, meaning subtle differences go undetected. We address this by developing a mathematical and statistical framework to study spheroid structure as a function of size, allowing us to ascertain whether initial spheroid size significantly affects growth dynamics.

Compared with traditional two-dimensional cell culture, spheroids closely mimic an avascular tumour microenvironment where spatial differences in the availability of nutrients strongly influence growth [14]. We observe that spheroids grow to a limiting size that is independent of the number of cells used to initiate the experiment (Fig. 1a–f), leading us to hypothesise that spheroids have a limiting structure [15]. This behaviour is consistent with untested predictions of mathematical models of tumour progression [16–25] (Fig. 1g). Many mathematical models assume that spheroid growth eventually ceases due to a balance between growth at the spheroid periphery and mass loss at the spheroid centre, driven by the spatial distribution of nutrients and metabolites (Fig. 1h) [16, 26]. We analyse highly detailed experimental data from a large number of spheroids to answer fundamental biological and theoretical questions. Firstly, we study the effect of initial spheroid size on the transient and limiting spheroid structure. The initial size of spheroids is often highly variable [14], yet is rarely accounted for in statistical analysis. Secondly, we study the relationship between spheroid size and structure using a mathematical model that describes growth inhibition due to the spatial distribution of nutrients and metabolites.

**Figure 1.**
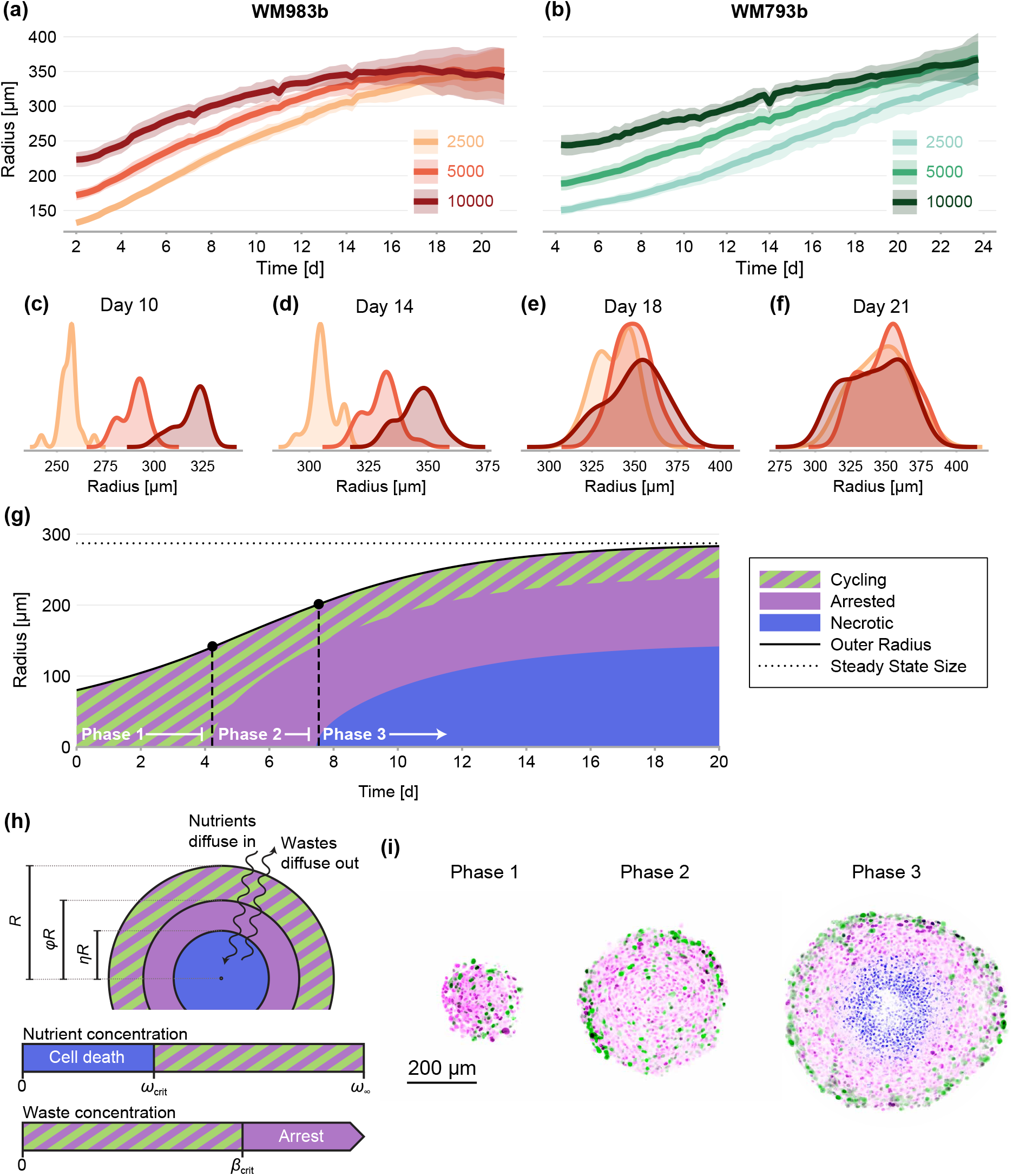
(a–f) Growth of WM983b and WM793b spheroids over three weeks, initiated using approximately 2500, 5000 and 10000 cells. The solid curve represents average outer radius and the coloured region corresponds to a 95% prediction interval (mean ± 1.96 std). (c–f) Size distribution of WM983b spheroids at day 10, 14, 18 and 21 for each initial seeding density. (g–h) Dynamics of the Greenspan model [16], which describes three phases of growth and the development of a stable spheroid structure under assumptions of nutrient and waste diffusion. We denote by *R* the spheroid radius, *ϕ* the relative radius of the arrested region and *η* the relative radius of the necrotic core. (i) Optical sections showing three phases of growth in the experimental data (WM983b spheroids initiated with 2500 cells at day 3, 7 and 14). Colouring indicates cell nuclei positive for mKO2 (magenta), which indicates cells in gap 1; cell nuclei positive for mAG (green), which indicates cells in gap 2; and cell nuclei stained with DRAQ7 (blue), which indicates necrosis.

We study spheroids grown at three seeding densities from human melanoma cells [27, 28] transduced with the fluorescent ubiquitination cell cycle indicator (FUCCI) [29–32]. FUCCI technology discriminates between cells in different stages of the cell cycle, namely gap 1 (before synthesis) and gap 2 (after DNA replication), allowing us to identify regions containing actively cycling cells, and regions where the majority of the cells are viable but in cell cycle arrest. We grow spheroids for up to 24 days to allow sufficient time to observe growth inhibition. We summarise experimental images using three measurements of spheroid structure: (1) the overall size of each spheroid; (2) the size of the inhibited region (which we define as the region where the majority of cells are in gap 1); and, (3) the size of the necrotic core.

It is widely accepted that the eventual inhibition of spheroid growth arises through three phases (Fig. 1g,i) [22,24,33]. During *phase 1*, for spheroids that are sufficiently small, we observe cycling cells throughout. In *phase 2*, spheroids develop to a size where cells in the spheroid centre remain viable but enter cell cycle arrest, potentially due to a higher concentration of metabolites in the spheroid centre [34, 35]. Finally, during *phase 3* the spheroid develops a necrotic core. Eventually, the loss of cells within the spheroid balances growth at the spheroid periphery, stalling net overall growth.

Whether spheroids reach the size required for necrosis to develop relates to experimental design choices such as the experimental duration and initial seeding density, among many other factors. Our hypothesis is that, provided the availability of nutrients is maintained in the cell culture, the structure of a spheroid is eventually a function of spheroid size, independent of the initial seeding density. This presents us with a technical challenge and a biological opportunity for protocol refinement. For example, we find that the initial aggregation of cells into spheroids occurs over several days [33], a timescale similar to that of cell proliferation. Therefore, the growth of spheroids over a short experimental duration may be significantly influenced by differences in initial seeding density, potentially confounding differences due to variations in cell behaviour between experimental conditions and limiting the reproducibility of experiments. Our analysis of late-time spheroid structure circumvents this by studying structure as function of overall size instead of time. The primary benefit of this approach is that inferences are insensitive to variations in the initial seeding density.

We take a likelihood-based approach to estimating parameters [36]; employ profile likelihood analysis to produce approximate confidence intervals [37–39]; and develop a likelihood-ratio-based hypothesis test to assess consistency in results between seeding densities. Firstly, we work solely with a statistical model that describes the average sizes of the spheroid, inhibited region and necrotic core at each observation time. Secondly, we apply a simple mechanistic model that describes spheroid progression due to a balance between growth at the spheroid periphery and mass loss due to necrosis in the spheroid centre. Following the seminal work of Greenspan [16], we assume that nutrients and wastes from living cells are at diffusive equilibrium, leading to a functional relationship between spheroid size and inner structure. Comparing model predictions to experimental observations allows us to assess whether the underlying assumptions of the Greenspan model are appropriate, providing valuable information for model refinement. As we are primarily interested in spheroid structure and model validation, we focus our analysis on comparing the structure at different observation times and seeding densities rather than a more typical approach that calibrates the mathematical to all data simultaneously [25].

We are motivated to work with a simple mathematical model instead of a more complex (and potentially more biologically realistic) alternative [20, 40–43] for two reasons. Firstly, complex models are often highly parameterised [44–46]. Given the practical difficulties in extracting detailed measurements from spheroids, we do not expect to be able to reliably estimate parameters in many complex models; that is, we expect parameters to be *practically non-identifiable* [37]. Working with a simple model avoids over-parameterisation allowing for a better comparison between experimental conditions. Secondly, Greenspan’s model encapsulates our central hypothesis that spheroid structure is purely a function of spheroid size, and captures the key features of spheroid growth seen in the experimental data with a low-dimensional, interpretable, parameter space.

## 2 Methods

### 2.1 Experimental methods

The human melanoma cell lines WM793b [27] and WM983b [28] were genotypically characterised [47–49], grown as described in [33] supplemented with 1% penicillin-streptomycin (ThermoFisher, Massachusetts, United States), and authenticated by short tandem repeat fingerprinting (QIMR Berghofer Medical Research Institute, Herston, Australia). All cell lines were transduced with fluorescent ubiquitination-based cell cycle indicator (FUCCI) constructs as described in [30, 33]. Wells within a flat-bottomed 96-well plate were prepared with 50 μL non-adherent 1.5% agarose to prevent cell-to-substrate attachment and promote the formation of a single centrally located spheroid [32]. Cells were seeded into each well at a density of approximately 2500, 5000 and 10000 cells in 200 μL of medium. A medium change was performed every 2 to 4 days.

Spheroids were harvested and fixed with 4% paraformaldehyde at day 3, 4, 5, 7, 10, 12, 14, 16, 18, 21 and 24; mounted in 2% low melting agarose; placed in a refractive-index-matched clearing solution [32]; and imaged using fluorescent confocal microscopy to obtain high-resolution images at the equator of each spheroid (Olympus FV3000, Olympus, Tokyo, Japan). To minimise variability due to the vertical position of each image, spheroids are fixed in place using an agarose gel, and equatorial images are defined as the cross-section with the largest cross-sectional area. To obtain the result in Fig. 1i, we selectively stain spheroids with DRAQ7™ (ThermoFisher, Massachusetts, United States), which indicates necrosis [31, 32]. Staining, fixation, and microscopy are repeated to obtain at least 20 WM983b spheroids at day 18 (spheroids initially seeded with 5000 and 10000 cells) and day 21 (spheroids seeded with 2500 cells); and at least 10 spheroids for all other conditions. Data are then randomly subsampled to obtain exactly 10 and 20 spheroids for each initial condition and observation day where possible. Time-lapse phase-contrast and fluorescent channel images are obtained at 6 hour intervals for up to 24 spheroids for each initial condition using an Incucyte S3 (Sartorius, Goettingen, Germany).

### 2.2 Data processing

We apply a semi-automated data processing algorithm to summarise experimental images with three measurements (Fig. 1h) [50]. Firstly, we calculate the outer radius, *R*, based on a sphere with the same cross-sectional area as the image obtained. Secondly, the radius of the inhibited region, *R*_i_. We calculate the radius of the inhibited region by determining the average distance from the spheroid periphery where the signal from mAG (FUCCI green), which indicates cells in gap 2, falls below a threshold value, taken to be 20% of the maximum area-averaged green signal. We find this choice leads to accurate results (Fig. 2). Finally, the radius of the necrotic core, *R*_n_, which is identified using texture recognition (`stdfilt`, [51]). The regions identified using the algorithm are shown in Fig. 2. Full details of the image processing algorithms are available in [50] and additional images are available as supplementary material.

**Figure 2.**
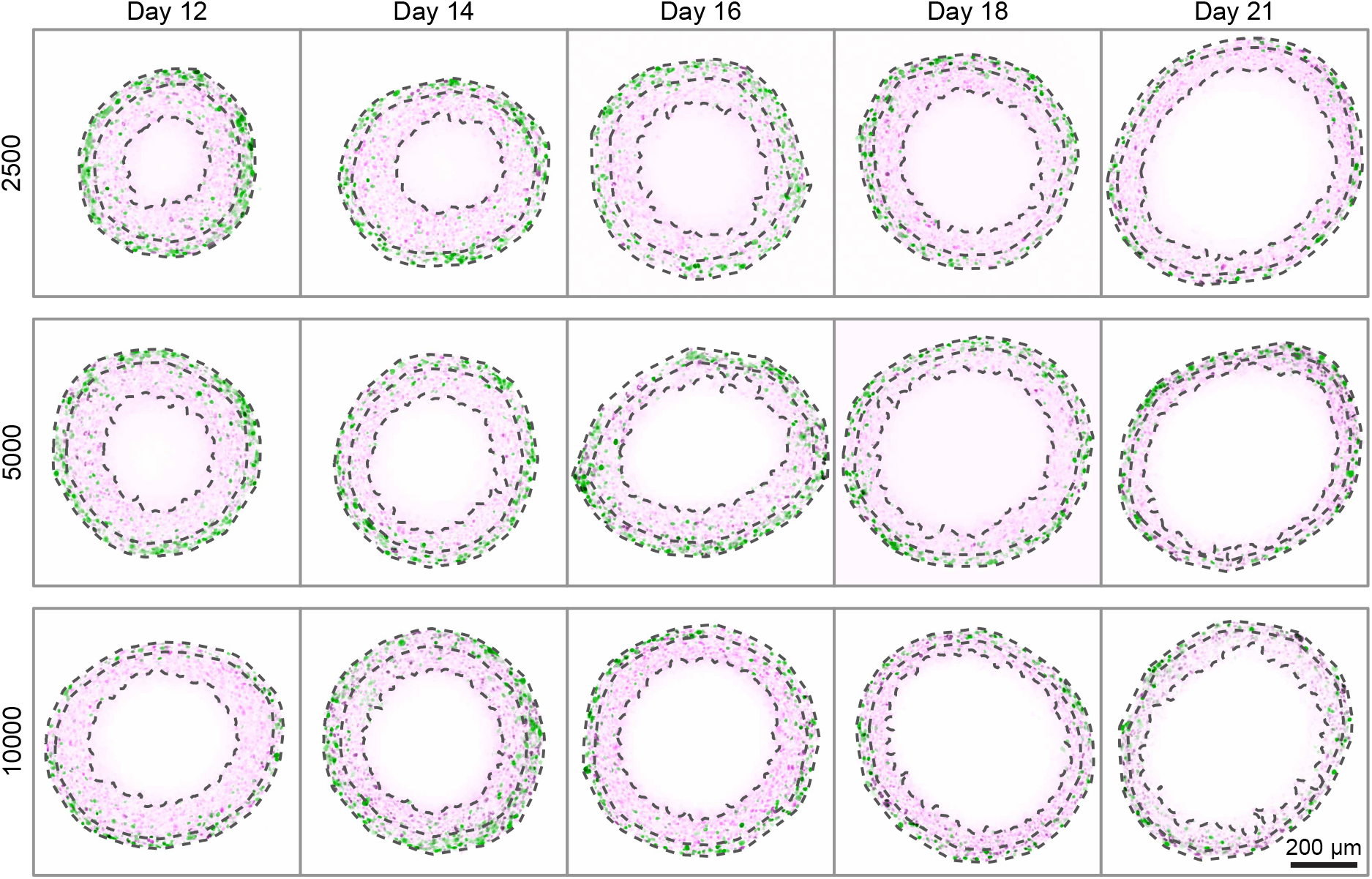
Late-time progression of WM983b spheroids, randomly sampled from the 10 spheroids imaged from each condition (additional images in Supplementary file 2). Overlaid are the three boundaries identified by the image processing algorithm: the entire spheroid, the inhibited region and the necrotic region. Each image shows a 800 × 800 μm field of view. Colouring indicates cell nuclei positive for mKO2 (magenta), which indicates cells in gap 1; and cell nuclei positive for mAG (green), which indicates cells in gap 2.

### 2.3 Mathematical model

Following [16], we make two minimal assumptions regarding growth inhibition and necrosis (Fig. 1h). Firstly, that growth inhibition, or cell cycle arrest, is a result of a chemical inhibitor that originates from the metabolic waste of living cells [52]. This inhibitor is produced by living cells at rate *β*_prod_ [mol d^−1^] and diffuses with diffusivity *β*_diff_ [μm^2^ d^−1^]. At the outer boundary of the spheroid, we assume that the concentration of inhibitor is zero. Cells enter arrest in regions where the inhibitor concentration is greater than *β*_crit_ [mol μm^−3^]. Secondly, cycling cells require nutrients that are plentifully available in the surrounding medium at concentration *ω*_∞_ [mol μm^−3^]. The nutrient is consumed by cycling cells at a constant rate *ω*_cons_ [mol d^−1^] and diffuses with diffusivity *ω*_diff_ [μm^2^ d^−1^]. Cells die in regions where the nutrient concentration is less than *ω*_crit_ [mol μm^−3^].

In regions where the nutrient concentration is sufficiently high and the inhibitor concentration sufficiently low, we assume that cells proliferate exponentially at the per-volume rate *s* [d^−1^]. Furthermore, we assume that cell debris is lost from the necrotic core at the per-volume rate *λ* [d^−1^].

It is convenient to define two non-dimensional parameters

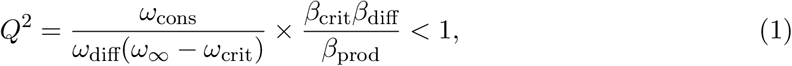

and

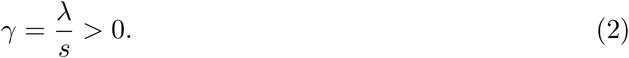

The parameter *Q* quantifies the balance between nutrient and inhibitor concentration and *γ* quantifies the balance between cell growth and the loss due to necrosis. The restriction *Q* < 1 arises since we observe an inhibited region form before the necrotic region [16]. Since the resultant equations depend only on *Q* and *γ*, the constituents of *Q*, namely *β*_prod_, *β*_diff_, *β*_crit_, *ω*_cons_, *ω*_diff_, *ω*_∞_ and *ω*_crit_, cannot be uniquely identified unless prior knowledge from other experiments is considered [53], perhaps in a Bayesian framework [54]. In contrast, the constituents of *γ*, namely *λ* and *s*, can be identified if information relating to the per-volume cell proliferation rate *s* is available, perhaps from phase 1 spheroid growth data.

We take the standard approach and model the spheroid as a single spherical mass [16, 21]. We denote by *R* the radius of the spheroid, *ϕ* = *R*_i_*/R* the relative radius of the inhibited region, and *η* = *R*_n_*/R* the relative radius of the necrotic core (Fig. 1h). We note that *R* > 0 and 0 ≤ *η* ≤ *ϕ* < 1. Noting that nutrient and inhibitor diffusion occurs much faster than cell proliferation, we assume that the chemical species are in diffusive equilibrium, leading to

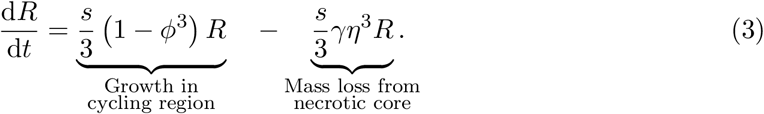

A distinguishing feature of Greenspan’s model is that the inner structure of the spheroid, quantified by (*ϕ, η*), is determined solely by the spheroid radius, and not by time. We denote

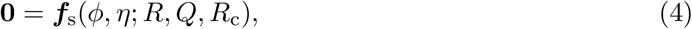

as a function describing this relationship, and refer to the relationship between the spheroid radius, *R*, and the inner structure, (*ϕ, η*), as the *structural model*. Here, we define *R*_c_ as the radius at which necrosis first occurs. For *R* < *R*_c_, nutrient is available throughout the spheroid above the critical concentration *ω*_crit_.

During phases 1 and 2, there is no necrotic core (*η* = 0) and the solution to Eq. 4 is given by

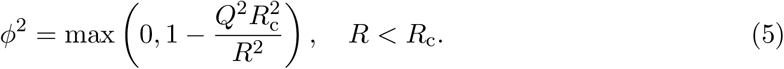

During phase 3, *R* > *R*_c_ and ***f***_s_ is given by

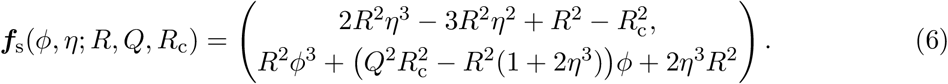

To investigate the limiting structure of spheroids, we consider the solution to the mathematical model where the outer radius is no longer increasing: the dynamics have reached a *steady-state*. Experimental observations suggest that this occurs during phase 3. We denote 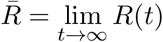 the limiting radius and 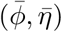 the associated limiting structure. The *steady-state model* is the solution of

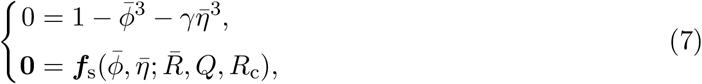

subject to *R* > *R*_c_. By defining 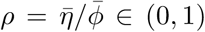, we find a semi-analytical solution to the steady-state model (Appendix 1).

The behaviour in the steady-state model is characterised by three parameters, ***θ*** = (*Q, R*_c_, *γ*). We denote the solution to Eq. 7 (i.e., the steady-state model) as

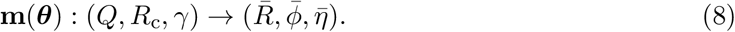

Equation 8 can be thought of as a map from the parameter space to the limiting structure of the spheroid. This demonstrates that the parameters are identifiable only when all three variables, 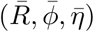, are observed, since the two-dimensional observation space (*R, η*) cannot uniquely map to the entire three-dimensional parameter space (*Q, R*_c_, *γ*). As a consequence, the model parameters cannot be uniquely identified from steady-state information unless phase 3 information that includes measurements of the inhibited region—using FUCCI or another marker of cell cycle arrest—is considered alongside measurements of necrotic core and overall spheroid size.

#### 2.3.1 Statistical model

While the mathematical model is deterministic, experimental observations of spheroid structure can be highly variable. To account for this, we take the standard approach and assume that the mathematical model describes the *expected behaviour* and experimental observations are multivariate normally distributed [36]. Aside from accounting for biological variability, the observation process captures variability introduced during imaging and image processing.

Denoting **x**_*i*_ = (*R*_*i*_, *ϕ*_*i*_, *η*_*i*_) as experimental observation *i* of the spheroid size and structure, we assume that

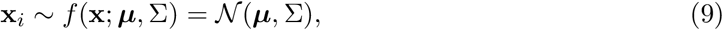

where ***μ*** = (*R, ϕ, η*) is the mean of each component of **x**, 𝒩(***μ***, Σ) denotes a multivariate normal distribution with mean ***μ*** and covariance Σ. To account for increased variability at later time points (Fig. 1a–b), we estimate Σ as the sample covariance associated with experimental observations of x_*i*_ at each time, *t*. For steady-state analysis, we calculate the covariance using the pooled sample covariance from all seeding densities.

We refer to Eq. 9 as the *statistical model*. To connect experimental observations to the *mathematical model*, we substitute ***μ*** = **m**(***θ***) in Eq. 9.

### 2.4 Inference

We take a likelihood-based approach to parameter inference and sensitivity analysis. Given a set of observations 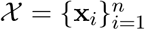, the log-likelihood function is

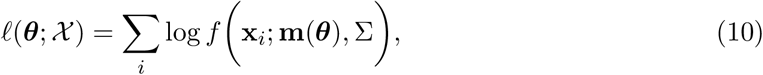

where *f* (**x**; ***μ***, Σ) is the multivariate normal probability density function (Eq. 9). Although we take a purely likelihood-based approach to inference, we note that our implementation is equivalent to a Bayesian approach where uniform priors encode existing knowledge about parameters, a common choice [55, 56].

We apply maximum likelihood estimation to obtain point estimates of the parameters for a given set of experimental observations. The maximum likelihood estimate (MLE) is given by

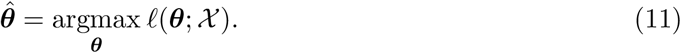

We solve Eq. 11 numerically to within machine precision using a local optimisation routine [57, 58]. In Fig. 3, we show point estimates obtained for a bivariate problem using maximum likelihood estimation.

**Figure 3.**
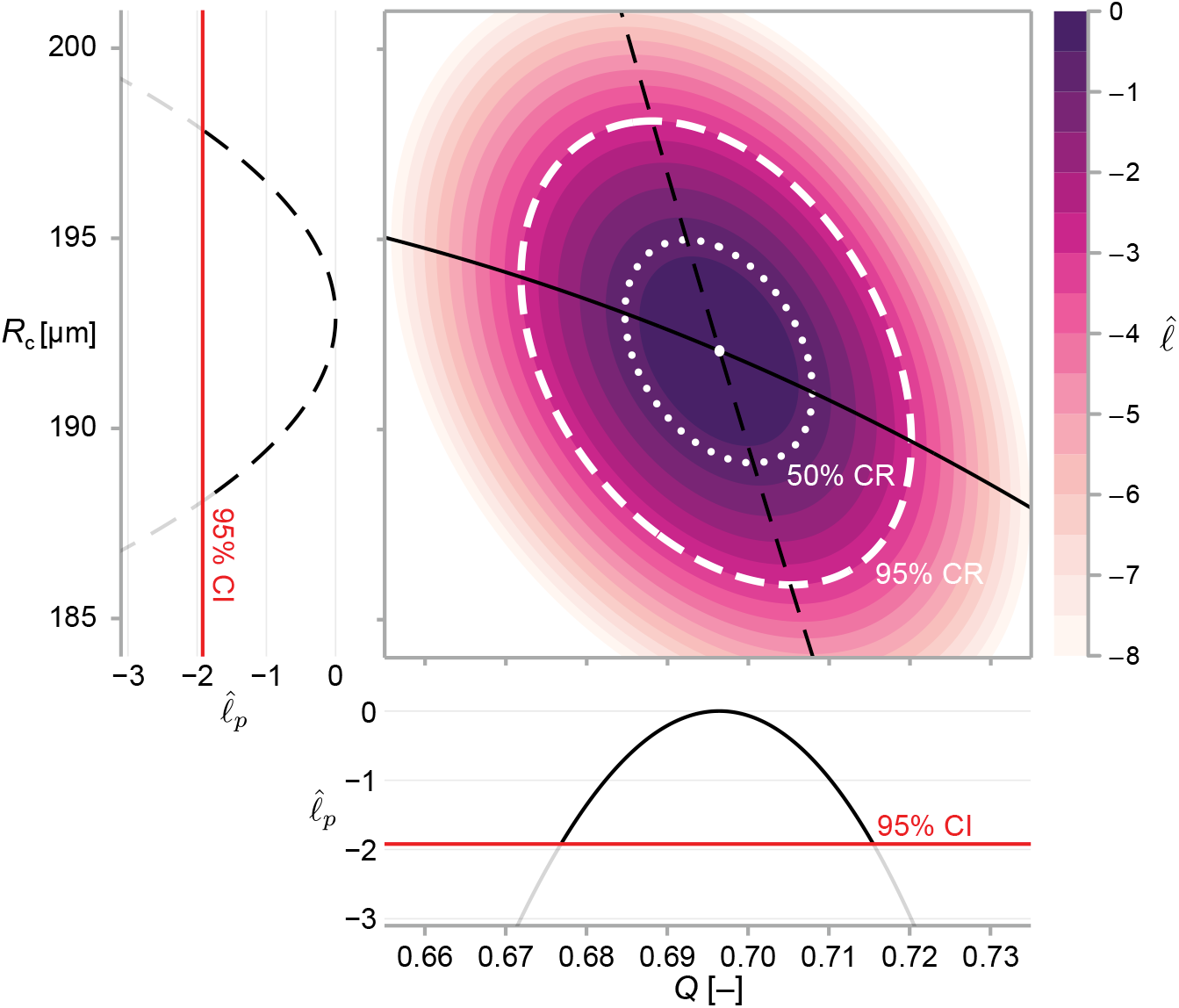
We calculate approximate confidence intervals (CI) using profile likelihood and confidence regions (CR) using contours of the normalised likelihood function. Results demonstrate estimates of *Q* and *R*_c_ using the structural model, Eq. 6, and data from WM983b spheroids at day 14 initiated using 5000 cells. Point estimates are calculated using the maximum likelihood estimate (white marker). The boundaries of regions are defined as contours of the log-likelihood function. Univariate confidence intervals are constructed by profiling the log-likelihood and using a threshold of approximately −1.92 for a 95% confidence interval.

#### 2.4.1 Confidence regions and hypothesis tests

We take a log-likelihood based approach to compute confidence regions and marginal univariate confidence intervals for model parameters [38]. In a large sample limit, Wilks’ Theorem provides a limiting distribution for the log-likelihood ratio statistic, such that

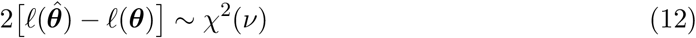

where *ν* = dim(***θ***) and *χ*^2^(*ν*) is the *χ*^2^ distribution with *ν* degrees of freedom. Therefore, an approximate *α* level confidence region is given by

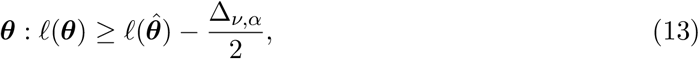

where Δ_*ν,α*_ is the *α* level quantile of the *χ*^2^(*ν*) distribution.

##### Hypothesis tests

To compare parameters between initial conditions, we perform likelihood-ratio-based hypothesis test based on the distribution provided in Eq. 13 [38]. We denote by 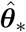 the MLE computed using data from all initial seeding densities, 𝒳_*_, simultaneously. Similarly, to compare parameter estimates from spheroids initially seeded with 2500 and 5000 cells, we denote by 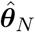 the MLE using a subset of data from spheroids seeded using *N* ∈ {2500, 5000} cells. The test statistic is given by

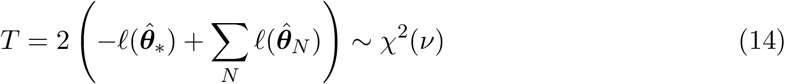

where *ν* is number of additional parameters in the case where a different parameter combination is used to describe each initial condition. An approximate *p*-value is therefore given by 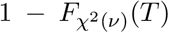, where 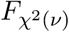 is the cumulative distribution function for the *χ*^2^(*ν*) distribution.

##### Marginal confidence intervals

The profile likelihood method [37,59] allows for the construction of univariate confidence interval of each parameter. Firstly, we partition the parameter space such that ***θ*** = (*ψ*, ***λ***) where *ψ* is the parameter of interest and ***λ*** is a vector containing the remaining parameters. Taking the supremum of the log-likelihood function over ***λ*** and normalising using the MLE gives the normalised profile log-likelihood

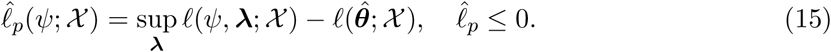

An approximate 95% confidence interval is given by Eq. 13 as the region where 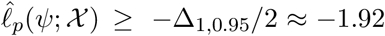 [38]. We compute the profile log-likelihood numerically using a local optimisation routine [57] with either the MLE, or the nearest profiled point [59] as an initial guess. In Fig. 3, we show profile likelihoods for a bivariate problem.

##### Confidence regions

We construct two-dimensional confidence regions using Eq. 13 (we construct three-dimensional confidence regions using a sequence of two-dimensional slices). First, we find a point on the boundary of the region, denoted ***θ***_0_ such that 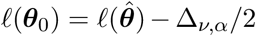, using bisection to machine precision. Next, we integrate along the *likelihood annihilating field* ; that is, we move in a direction perpendicular to the gradient of the likelihood to obtain a set of points on the level set 𝓁(***θ***) = 𝓁(***θ***_0_), given by

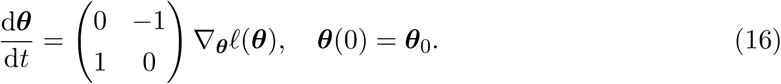

This calculation is demonstrated for a bivariate problem in Fig. 3.

The gradient for the statistical model, ∇_***μ***_𝓁(***μ***), can be calculated to within machine precision using automatic differentiation [60]. For the mathematical model, we apply the identity

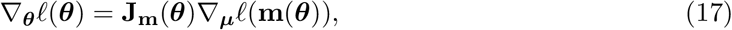

where **J**_**m**_(***θ***) is calculated analytically (Appendix 1).

## 3 Results

To assess the limiting structure of spheroids and the effect of initial seeding density, we analyse confocal sections of a large number of spheroids across three seeding densities using the WM983b cell line. We show a subset of these images in Fig. 2 and summarise images with three concentric annular measurements: the spheroid radius, *R*; the relative radius of the inhibited region, *ϕ*; and the relative radius of the necrotic core, *η* (Fig. 1h). In addition to spheroids from different initial conditions tending towards a similar overall size (as seen from time-lapse data in Fig. 1a–f), these results show that spheroids develop similar structures by day 21.

First, we fit the statistical model to the experimental data by estimating the mean of each measurement, denoted ***μ*** = (*R, ϕ, η*). We obtain a maximum likelihood estimate and an approximate 95% confidence interval for each initial condition at observation days 12 to 21 (Fig. 4a–c). On average, spheroids of all seeding densities increase in size from day 12 to day 18. In agreement with earlier observations from time-lapse data in Fig. 1e–h, we see that spheroids initiated at different seeding densities tend toward similar limiting sizes. Between days 18 to 21, spheroids seeded with 5000 and 10000 cells decrease in average size, potentially indicating a period of decay after a limiting size is reached.

**Figure 4.**
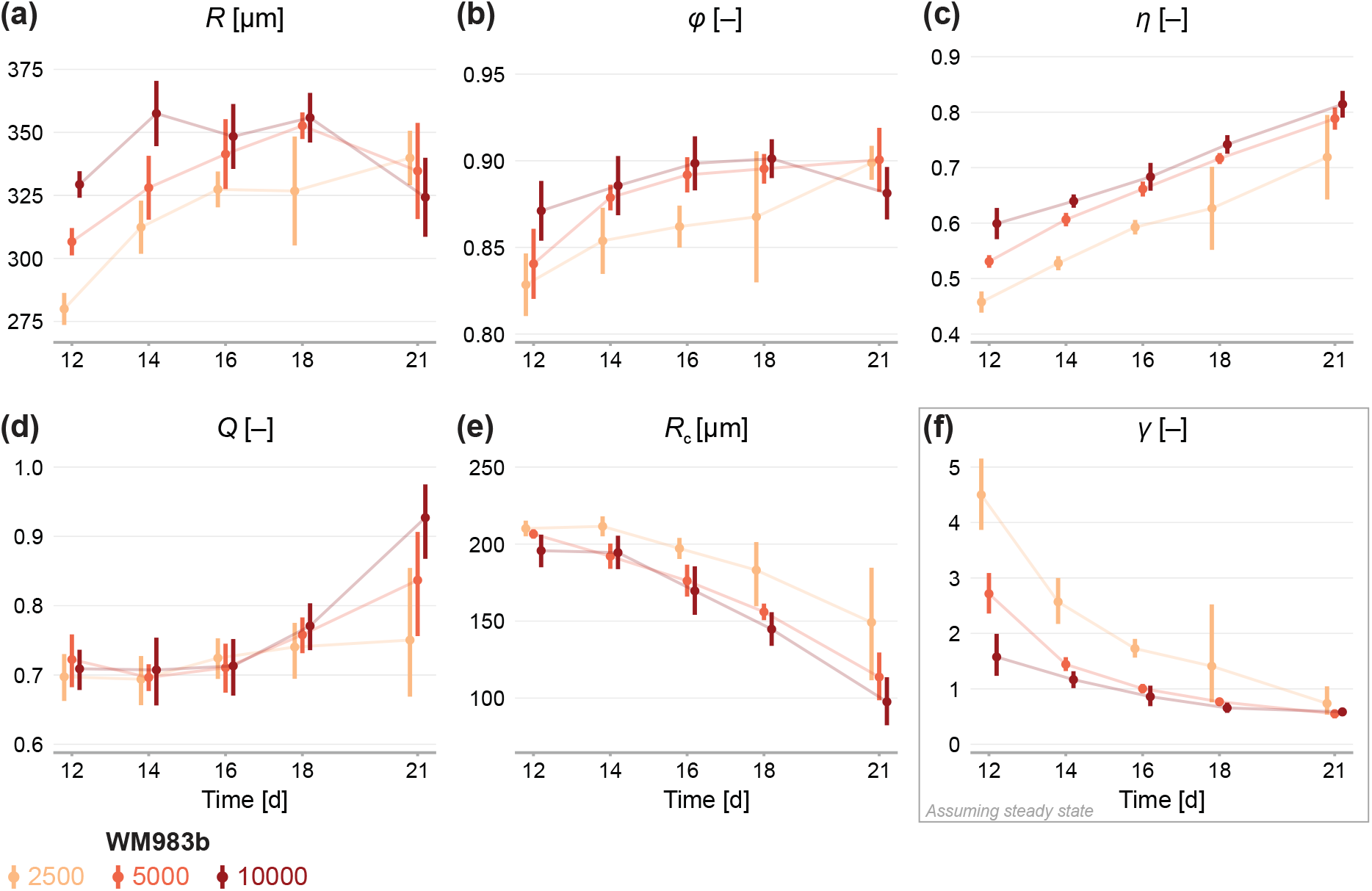
Estimates of parameters using the structural model with data from various time points. In (a–c), parameters are the mean of each observation: (*R, ϕ, η*). In (d–e), parameters are those in the structural model: (*R, Q, R*_c_). In (f), estimates of *γ* are obtained by calibrating observations to the steady-state model. As estimates *Q* and *R*_c_ can be derived from the structural model (Eq. 6), which applies at any time during phase 3, we expect to see similar parameter estimates across observation times. As estimates of *γ* can only be obtained from the steady-state model, which assumes the outer radius is no longer increasing, we do not expect to see similar parameter estimates across observation times. Bars indicate an approximate 95% confidence interval.

Figure 4b and c show estimates relating to the sizes of the inhibited region, *ϕ*, and necrotic core, *η*. We see remarkable consistency in *ϕ* across seeding densities, tending toward a value of 90% in all cases: this corresponds to an actively cycling region with volume approximately 27% of the total spheroid volume. The necrotic core increases significantly in size from days 12 to 21, and late time estimates of *η* are quantitively similar between seeding densities.

Next, we calibrate the mathematical model to identify any mechanistic differences between seeding densities. Parameters *Q* and *R*_c_ can be estimated using the structural model (Eq. 6) at any time point. To estimate *γ* we must invoke the steady-state model (Eq. 7), which assumes that the overall growth of the spheroid has ceased. Therefore, we expect to see consistency in estimates of *Q* and *R*_c_ between observation days but do not expect the same for estimates of *γ*.

Results in Fig. 4d show remarkable consistency in estimates of *Q* across seeding densities until day 18, suggesting that the balance between nutrient availability and waste concentration (Eq. 1) is maintained throughout the experiment and is similar between seeding densities. Between days 18 and 21, estimates of *Q* for spheroids initially seeded with 5000 and 10000 cells increase significantly, suggesting a behavioural change during this time; we attribute this to a final period of decay. Estimates of *R*_c_ do not show the consistency between observation days we might expect if **f**_s_ (Eq. 6) holds for the experimental data. Rather, estimates of *R*_c_ decrease between days 12 to 21, indicating **f**_s_ may be misspecified. Results in Fig. 4f show that estimates of *γ* decrease with time to a similar value for all seeding densities. We interpret this asymptotic decrease as an indication that spheroids approach a limiting structure since estimates of *γ* are strictly only valid when growth has ceased. Closer inspection of results in Fig. 4f show a delay in estimates of *γ* between spheroids seeded with 2500 cells and the other seeding densities. Whereas the larger spheroids reach a limiting size by day 18, the smaller spheroids are still growing. It is not until day 21 that estimates of *γ* are comparable across all seeding densities.

Next, we analyse the limiting structure of spheroids across each initial seeding density. As spheroids initially seeded with 5000 and 10000 cells decrease in average size from day 18 to day 21, we compare day 18 data from these high densities to day 21 data from spheroids initially seeded with 2500 cells. Results in Fig. 5a–c show profile log-likelihoods for each parameter in the mathematical model. In Fig. 5d and e, we show 3D confidence regions for parameters in the statistical and mathematical models, respectively. We see that both profile log-likelihoods and 3D confidence regions overlap, indicating that parameter estimates are consistent between seeding densities.

**Figure 5.**
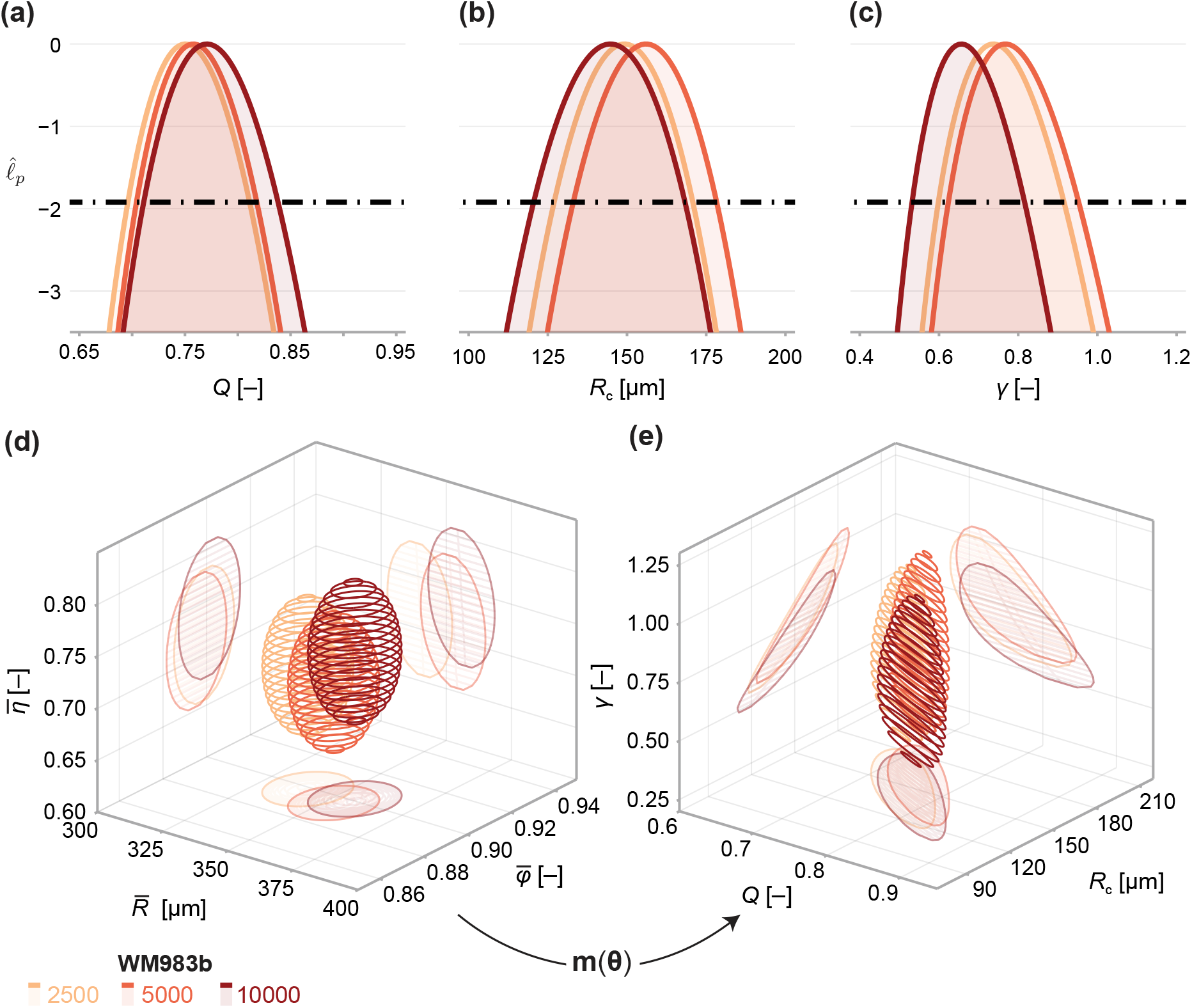
Comparison of WM983b spheroids between each initial seeding density at day 18 (spheroids seeded with 5000 or 10000 cells) and day 21 (2500). (a–c) Profile likelihoods for each parameter, which are used to compute approximate confidence intervals (Table 1). (d) 95% confidence region for the full parameter space. 95% confidence regions for (d) the mean of each observation at steady state 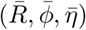 and (e) the model parameters (*Q, R*_c_, *γ*).

To compare quantitatively parameter estimates between seeding densities, we tabulate maximum likelihood estimates, approximate 95% confidence intervals, and results of a likelihood-ratio-based hypothesis test for both models in Table 1. The late-time sizes of spheroids initiated with 5000 and 10000 cells are statistically consistent (*p* = 0.62), as is their structure (*p* = 0.69). We find evidence to suggest that spheroids seeded with 2500 cells, even at day 21, are smaller (*p* = 0.04); however, the overall size and structure of the spheroids seeded with 2500 and 5000 cells are statistically consistent (*p* = 0.20). We find no significant differences in model parameters between seeding densities and note that the conclusion of overall statistical consistency between seeding densities is identical for the mathematical model.

**Table 1.** Parameter estimates and approximate confidence intervals for each initial conditions. Also shown are *p*-values for likelihood-ratio-based hypothesis tests for equivalence between seeding densities.

| Parameter | $\theta_{2500}$ | | $\theta_{5000}$ | | $\theta_{10000}$ | | $p_{2500,5000}$ | $p_{5000,10000}$ |
| --- | --- | --- | --- | --- | --- | --- | --- | --- |
| $R$ | 340.0 | (331.0, 349.0) | 353.0 | (344.0, 361.0) | 356.0 | (347.0, 365.0) | 0.0420 | 0.617 |
| $\phi$ | 0.899 | (0.889, 0.908) | 0.895 | (0.886, 0.905) | 0.901 | (0.891, 0.911) | 0.617 | 0.406 |
| $\eta$ | 0.719 | (0.674, 0.764) | 0.716 | (0.671, 0.761) | 0.742 | (0.696, 0.788) | 0.940 | 0.438 |
| $\mu$ | | | | | | | 0.202 | 0.687 |
| $Q$ | 0.75 | (0.696, 0.811) | 0.758 | (0.704, 0.818) | 0.771 | (0.711, 0.838) | 0.854 | 0.767 |
| $R_{\text{crit}}$ | 149.0 | (127.0, 171.0) | 156.0 | (133.0, 178.0) | 145.0 | (121.0, 168.0) | 0.672 | 0.503 |
| $\gamma$ | 0.737 | (0.598, 0.916) | 0.768 | (0.624, 0.953) | 0.657 | (0.532, 0.816) | 0.792 | 0.308 |
| $\theta$ | | | | | | | 0.202 | 0.687 |

Next, we investigate the relationship between spheroid structure and spheroid size from day 3 to day 21 (Fig. 6a). We again see evidence of a period of eventual decay that occurs after a limiting size has been reached in our experiments. To validate the structural relationship suggested by Greenspan’s model, we plot the solution to the structural model (Eq. 6) using parameters estimated using the steady-state model (Table 1). The overall trend throughout all three phases of growth in the mathematical model—made only using information from days 18 and 21—is remarkably consistent with experimental measurements Fig. 6a. We find an explanation for the inconsistent estimates of *R*_c_ observed in Fig. 4e. During phase 3, the mathematical model predicts a non-linear relationship between *R, ϕ* and *η* (Eq. 6). In contrast, the trend in the data is close to linear. We confirm this in Fig. 6b by calibrating a linear model of the form

**Figure 6.**
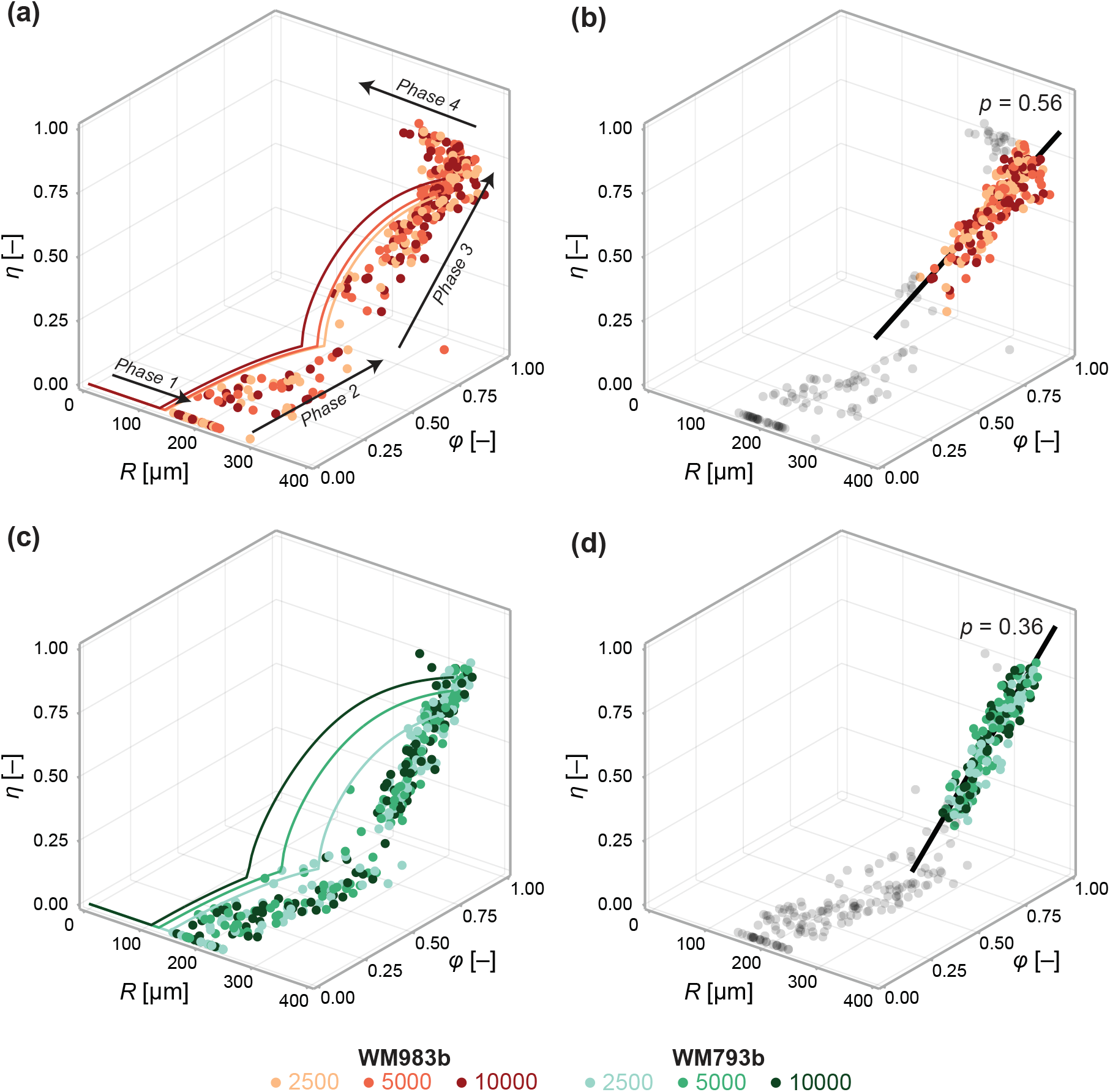
Data from days 3 to 21 (WM983b) and days 4 to 24 (WM793b) for all initial conditions. Solid curves in (a) show the solution to the mathematical model (Eq. 6) using the maximum likelihood estimate calculated using the steady-state data (Table 1). Solid curves in (c) show the solution to the mathematical model (Eq. 6) using using the maximum likelihood estimate calculated using day 24 data. In (b) and (d), we fit a linear model to phase 3 data (indicated by coloured markers). The *p* value corresponds to a hypothesis test where the linear model parameters are the equivalent for all initial conditions. Shown in black is the best fit linear model.

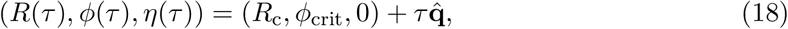

to phase 3 data using a total least squares approach that accounts for uncertainty in the independent variable *τ* (Appendix 2). Here, *τ* = 0 at the start of phase 3. Performing an approximate likelihood-ratio-based hypothesis test confirms that the behaviour in spheroids of all initial conditions is statistically consistent (*p* = 0.56). That is, the spheroid structure where necrosis first occurs (at *τ* = 0), (*R*_c_, *ϕ*_crit_, 0), and the direction in which it develops, 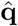, do not appear to depend on the initial seeding density.

In Fig. 6c and 6d, we perform a similar analysis on spheroids grown from WM793b cells. Whereas WM983b spheroids approach a limiting size by the conclusion of the experiment (Fig. 1a), spheroids grown from the WM793b do not (Fig. 1b). Results in Appendix 3 examine parameter estimates from the mathematical and statistical models through time for the WM793b spheroids, demonstrating that the outer radius increases monotonically until day 24 for all initial conditions. These results also suggest consistency in estimates of *Q* across observation days and seeding densities. Performing a likelihood-ratio-based hypothesis test indicates that phase 3 is independent of the initial seeding density (*p* = 0.36).

## 4 Discussion

Time-lapse measurements of WM983b spheroids over a 21-day experiment show a cessation in overall growth as the spheroids reach a limiting size. Consistent with largely untested predictions of classical mathematical models [16,20,21], these limiting sizes appear to be independent of the initial seeding density. Motivated by these observations, we develop a quantitative framework to study spheroid structure as a function of overall size. We aim to answer two fundamental questions: Do these spheroids have a limiting structure? Is the late-time behaviour independent of the initial seeding density?

We find compelling evidence that WM983b spheroids have a limiting structure that is independent of the initial seeding density. This assumption is routinely invoked in mathematical models of tumour structure but is yet to be experimentally verified. Given that we observe spheroids to eventually reduce in size, we compare structural measurements at days when the average outer radius for each initial seeding density is largest. First, we establish that spheroids seeded with 5000 and 10000 cells have similar limiting sizes (353 μm and 356 μm, respectively; *p* = 0.62) and that spheroids seeded with 2500 cells are slightly smaller at late time (340 μm; *p* = 0.04). This result highlights one of the challenges in determining the limiting structure of spheroids: it is unclear whether there is a difference or whether the smaller spheroids would continue to grow given additional time. Despite this discrepancy, we find a statistically consistent limiting structure, with a necrotic core of 73% of the outer radius and an inhibited region of 90% of the outer radius, indicating a proliferative periphery approximately 35 μm (two to three cell diameters) thick.

By examining spheroid structure throughout the entire experiment (Fig. 6), we establish a relationship between spheroid structure and size that is independent of initial seeding density. This result is significant as it suggests that variability in size and structure may be primarily attributable to time. For example, spheroids that are smaller than average on a given observation day may have been seeded at a lower density. Statistical techniques, such as ODE-constrained mixed effects models, can be applied to elucidate sources of intrinsic variability, such as variability in the initial seeding density [61,62]. It is common in the literature to compare spheroids with and without a putative drug after a fixed number of days [12]. However, our analysis suggests that comparing the structure of spheroids of a fixed size may be more insightful; this approach obviates variability due to initial seeding density, increasing the sensitivity of statistical tests to small effects. A corollary is that since inferences relating to spheroid structure are independent of spheroid size, experiments can be initiated with a larger number of cells to decrease the time until spheroids reach phase 3.

Given our observations of WM983b spheroids across seeding densities, an apparent conclusion of our analysis is that statistically consistent phase 3 behaviour implies a statistically consistent limiting structure. If true, this suggests that an experimentalist only has to investigate phase 3 behaviour to reach a conclusion relating to the limiting structure. Analysis of both the mathematical model and experimental results for WM793b spheroids indicate that this is not the case. In the mathematical model, ***f***_s_ (Eq. 6) characterises the structure solely in terms of parameters *Q* and *R*_c_, whereas *γ*—which relates to the ratio of cell proliferation and loss due to necrosis (Eq. 2)—determines the steady-state. We see this for WM793b spheroids, as phase 3 behaviour is independent of the initial seeding density (Fig. 6d), but time-lapse data of the overall growth (Fig. 1b) gives no indication that spheroids of different densities will tend toward the same limiting size.

As ***f***_s_ determines the relationship between spheroid size and structure at any time point, we expect estimates of *Q* and *R*_c_ to be similar when calibrated to data from different days. This is the case for estimates of *Q* (Fig. 4d), but estimates of *R*_c_ decrease with time (Fig. 4e). While the mathematical model captures the same overall behaviour observed in the experiments, it is evident from the discrepancy observed during phase 3 (Fig. 6a) that ***f***_s_ is misspecified. Our assumptions of nutrient and waste at diffusive equilibrium and a hard threshold for growth inhibition and necrosis give rise to ***f***_s_ that is cubic in *ϕ* and *η*. Since the empirical relationship for the cell lines we investigate is approximately linear, the model underestimates the radius at which phase 3 begins, *R*_c_. At the loss of mechanistic insight, one approach to rectify this discrepancy is to construct a purely phenomenological relationship where ***f***_s_ is piecewise linear. A second approach is to revisit fundamental modelling assumptions to develop a mechanistic description of the relationship between spheroid structure and overall size that is consistent with our experimental observations for these cell lines.

Our observations for WM983b and WM793b melanoma cell lines do not preclude a form of ***f***_s_ that is cubic for other cell lines or experimental conditions. In our framework, the behaviour of spheroids is characterised by the empirical relationship between spheroid size and structure. Therefore, despite misspecification in parameter estimates of *R*_c_, we can compare spheroids grown with WM793b and WM983b cell lines by comparing the structural relationship observed in the experimental data (Fig. 6a and 6c). In this case, we observe that radius at which the necrotic core develops is much smaller in WM983b spheroids than for WM793b spheroids. While we cannot elucidate the biological factors that lead to this difference from our analysis, we postulate that differences in the diffusion or consumption of nutrients by cells of each cell line may contribute.

We have restricted our analysis of spheroid structure to three measurements that quantify the sizes of the spheroid, inhibited region and necrotic core. While the spheroid and necrotic core sizes are objective measurements, the boundary of the inhibited region is not. Our approach is to identify the distance from the spheroid periphery where the density of cells in gap 2 falls below 20% of the maximum. We find this semi-automated approach produces excellent results and enables high-throughput analysis of hundreds of spheroids; however, it does not take advantage of all the information available in the experimental images. Mathematical models that explicitly include variation in cell density through space [20, 63] may be appropriate, however are typically heavily parameterised, limiting the insight obtainable from typical experimental data. The mass-balance model coupled to a model describing the relationship between spheroid size and structure avoids these issues and, despite model simplicity, we are still able to gain useful biological insight.

## 5 Conclusion

Reproducibility and size uniformity are paramount in practical applications of spheroid models. Yet, the effect of intentional or unintentional variability in spheroid size on the inner structure that develops is not well understood. We present a quantitative framework to analyse spheroid structure as a function of overall size, finding that the outer radius characterises the inner structure of spheroids grown from two melanoma cell lines. Further, we find that the initial seeding density has little effect on the structure that develops. These results attest to the reproducibility of spheroids as an *in vitro* research tool. While we analyse data from two melanoma cell lines, our focus on commonly reported spheroid measurements allows our framework to be applied more generally to a other cell lines and culture conditions. It is routine to compare spheroid size and structure of spheroids at a pre-determined time, our results suggest a refined protocol that compares the structure of spheroids at a pre-determined overall size.

Given the prominence of spheroids in experimental research, there is a surprising scarcity of experimentally validated mathematical models that can be applied to interpret data from these experiments. We find that one of the earliest and simplest models of tumour progression— the seminal model of Greenspan [16]—can give valuable insights with a parameter space that matches the level of detail available from spheroid structure data. Given that we establish an empirical relationship between spheroid size and structure independent of both time and the initial spheroid size, we suggest future theoretical work to identify mechanisms that give rise to this relationship, perhaps through equation learning [64]. To aid in validating theoretical models of spheroid growth, we make our highly detailed experimental data freely available.

## Data availability

Code, data, and interactive figures are available as a Julia module on GitHub at github.com/ap-browning/Spheroids. Code used to process the experimental images is available on Zenodo [50].

## Additional files

- Supplementary file 1. Spheroid count per experimental condition (harvest day, seeding density and cell line).
- Supplementary file 2. Additional cross-sectional confocal images of spheroids; 10 per experimental condition (harvest day, seeding density and cell line).
- Supplementary file 3. Reproduction of Fig. 5 using data from day 21 for all initial seeding densities.

## Acknowledgements

We thank Patrick Thomas for helpful comments and discussions and John Blake for guidance using the Incucyte S3. We thank Jennifer Flegg and one anonymous referee for helpful comments.

## Author contributions

All authors conceived the study. A.P.B. drafted the manuscript. A.P.B. and J.A.S. performed the analysis. A.P.B., R.J.M. and G.G. performed the experiments. A.P.B. and R.J.M. processed the experimental data. All authors provided comments and gave approval for publication.

## Appendix 1

### Steady-state model solution

The steady-state, denoted 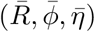 is given by setting d*R/* d*t* = 0 (Eq. 3 in the main text) yielding the non-linear system of equations

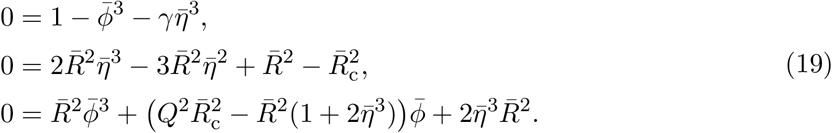

Applying the substitution 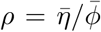, where 0 ≤ *ρ* ≤ 1, and algebraic manipulation allows the solution to Eqs. 19 to be expressed as the root of *f* (*ρ*; *Q, γ*), where

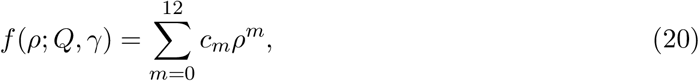

and where

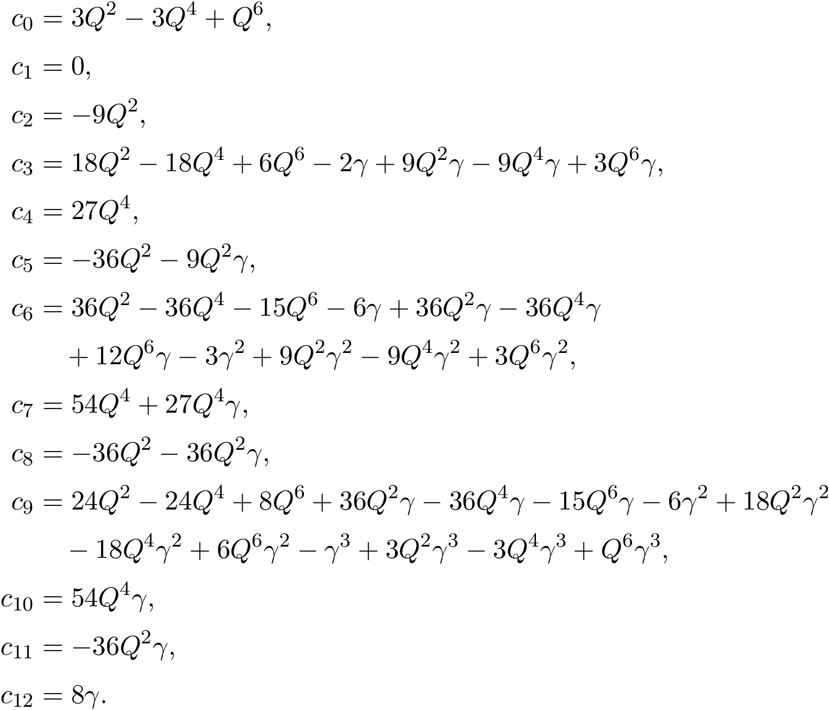

Since *ρ* is subject to the constraint 0 ≤ *ρ* ≤ 1, we solve 0 = *f* (*ρ*; *Q, γ*) using bisection^1^, which is guaranteed to converge provided there exists only one root in the interval 0 ≤ *ρ* ≤ 1. In Fig. A1a, we demonstrate that in the parameter region of interest (0 < *Q* < 1, *γ* > 0) there exists only a single solution to Eq. 20. We do this by finding all 12 roots of Eq. 20^2^ and counting the number of real roots where 0 ≤ *ρ* ≤ 1.

The solution to Eq. 19 is then given by

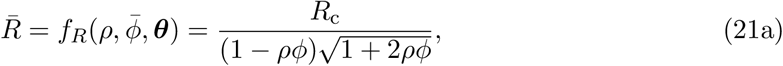

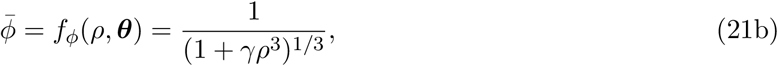

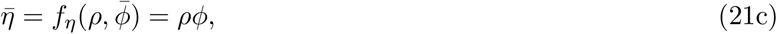

where ***θ*** = (*Q, R*_c_, *γ*).

In Fig. A1b, we compare a numerical solution to the transient model to the semi-analytical solution for the steady state showing an excellent match. All algorithms used to produce the results relating to the mathematical model are available on Github in `Module/Greenspan.jl`.

**Figure A1.**
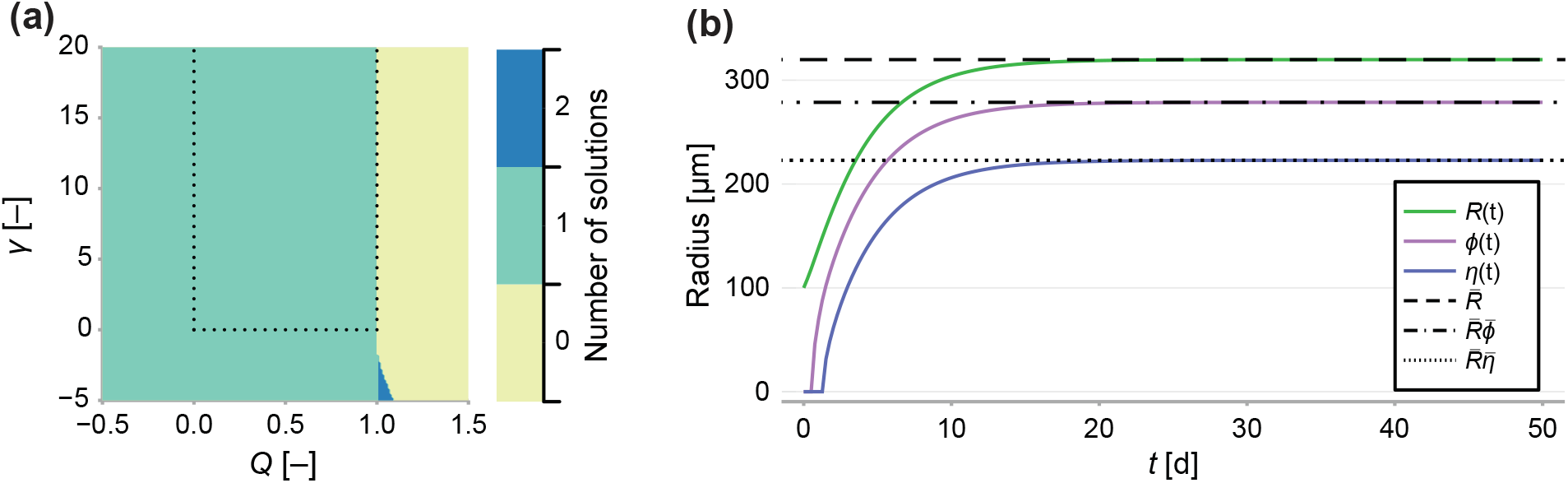
(a) Number of solutions to Eq. 20 subject to the constraint 0 ≤ *ρ* ≤ 1. Dashed line indicates the region of interest, where *γ* > 0 and 0 < *Q* < 1. (b) Comparison between a long-term solution to the transient model and the semi-analytical solution to the steady state, where *Q* = 0.8, *γ* = 1, *R*_c_ = 150, *s* = 1 and *R*_0_ = 100.

### Jacobian of the steady-state model

In the main document, we denote the solution to Eq. 19 as **m**(***θ***). Here, we demonstrate how given a value 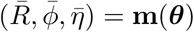, we can obtain an analytical expression for the model Jacobian,

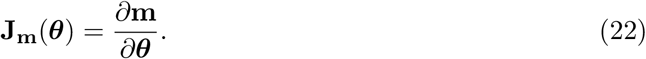

Given *ρ*, we can form an analytical expression for Eq. 22. Noting that the coefficients of Eq. 20 are functions of ***θ***, we consider

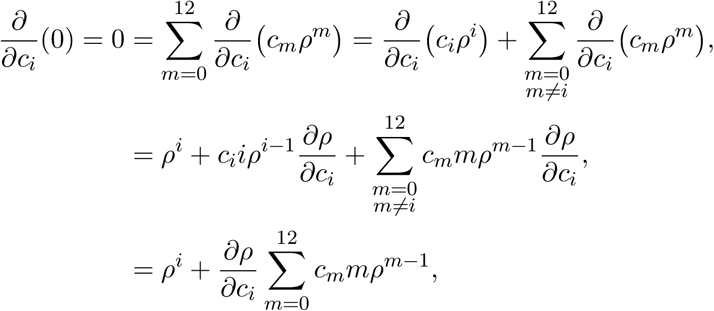

which yields

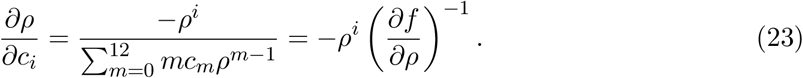

Therefore,

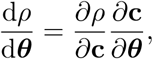

where **c** = (*c*_0_, *c*_1_, …, *c*_12_); *∂ρ/∂***c** = (*∂ρ/∂c*_0_, …, *∂ρ/∂c*_12_) and *∂***c***/∂****θ*** is the Jacobian of **c** with respect to ***θ***.

Therefore, we have that

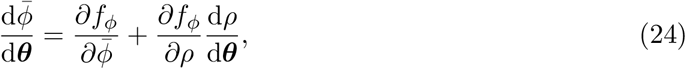

and it follows that

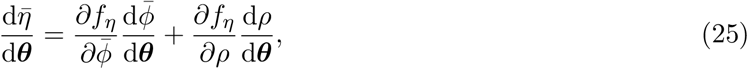

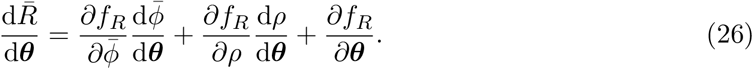

Therefore, an analytical expression for **J**_**m**_(***θ***) (Eq. 22) is given by

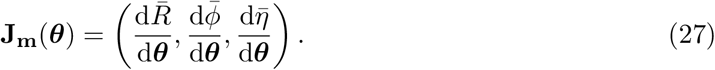

## Appendix 2

### Total squares regression

In typical least-squares estimation we fit a model of the form

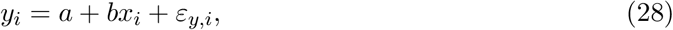

where *ε*_*y,i*_ ∼ 𝒩 (0, *σ*_*y*_) is assumed to be a normally distributed error component in *y* component [65], and (*a, b*) are model parameters. Least-squares and maximum likelihood estimates 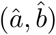 can then be found by minimising the sum-square error

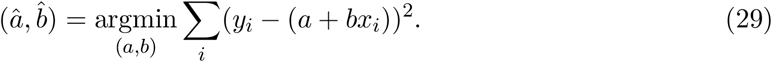

We demonstrate this in Fig. A3. In typical least squares estimation, we minimise the *vertical distance* between the data points and the regression line (blue dashed).

In the main document, we fit a linear model to data of the form (*R, ϕ, η*), where each component contains an error term. In two-dimensions, this is akin to a model of the form

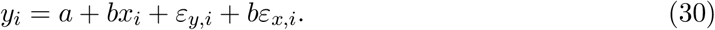

where we have included an additional error term *ε*_*x,i*_ ∼ 𝒩 (0, *σ*_*x*_), assumed to be a normally distributed error component in *x*_*i*_. In this case, the least squares estimate is given by minimising the total *perpendicular distance* between the data points and the regression line (Fig. A3, blue solid) [65].

In the main paper, we fit a linear model of the form

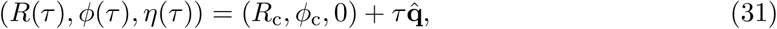

parameterised by *R*_c_, *ϕ*_c_ and a unit vector 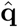.

If we denote y_0_ = (*R*_c_, *ϕ*_c_, 0) and 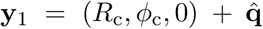, then the shortest distance between observation x_*i*_ = (*R*_*i*_, *ϕ*_*i*_, *η*_*i*_) is given by

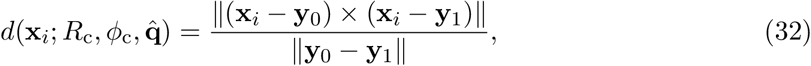

where ∥ · ∥ denotes the Frobenius norm, and × denotes the vector cross product.

Therefore, least-squares estimates of the parameters can then be found by minimising the sum-square error

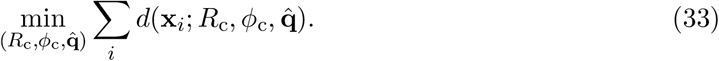

### Approximating the likelihood

To implement a log-likelihood-ratio based hypothesis test, we must approximate the likelihood at the parameter estimates. To do this, we note that the total square error, denoted 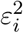, is of the form

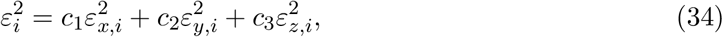

where *ε*_*x,i*_, *ε*_*y,i*_, and *ε*_*z,i*_ are normally distributed with variances 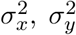 and 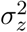, respectively. If 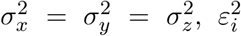 would have an approximate chi-squared distribution by the Welch-Satterthwaite equation [66], a special case of the gamma distribution. Therefore, we approximate the distribution of 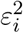 by fitting a gamma-distribution to the observed square error when a total squares estimate is fit to the combined data (Fig. A3b).

Therefore, the approximate log-likelihood is given by

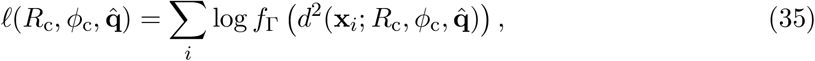

where *f*_Γ_(·) is the probability density function of the fitted gamma function.

### Log-likelihood-ratio based test

We denote 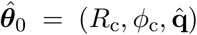 the maximum likelihood estimate when the data from all initial conditions is pooled, and 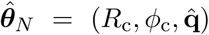 the estimates from initial condition *N* ∈ {2500, 5000, 10000}. As the models must be nested for the likelihood-ratio test, we estimate the noise model, *f*_Γ_(·), using the estimates from the pooled data.

The test-statistic is given by

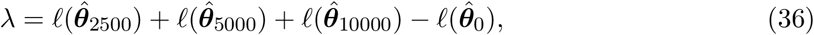

where 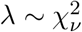, and

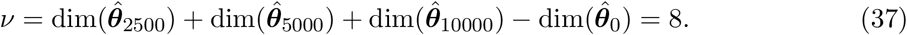

Our implementation of this test is provided on GitHub in `Module/Inference` in the function `lm orthogonal test`.

**Figure A2.**
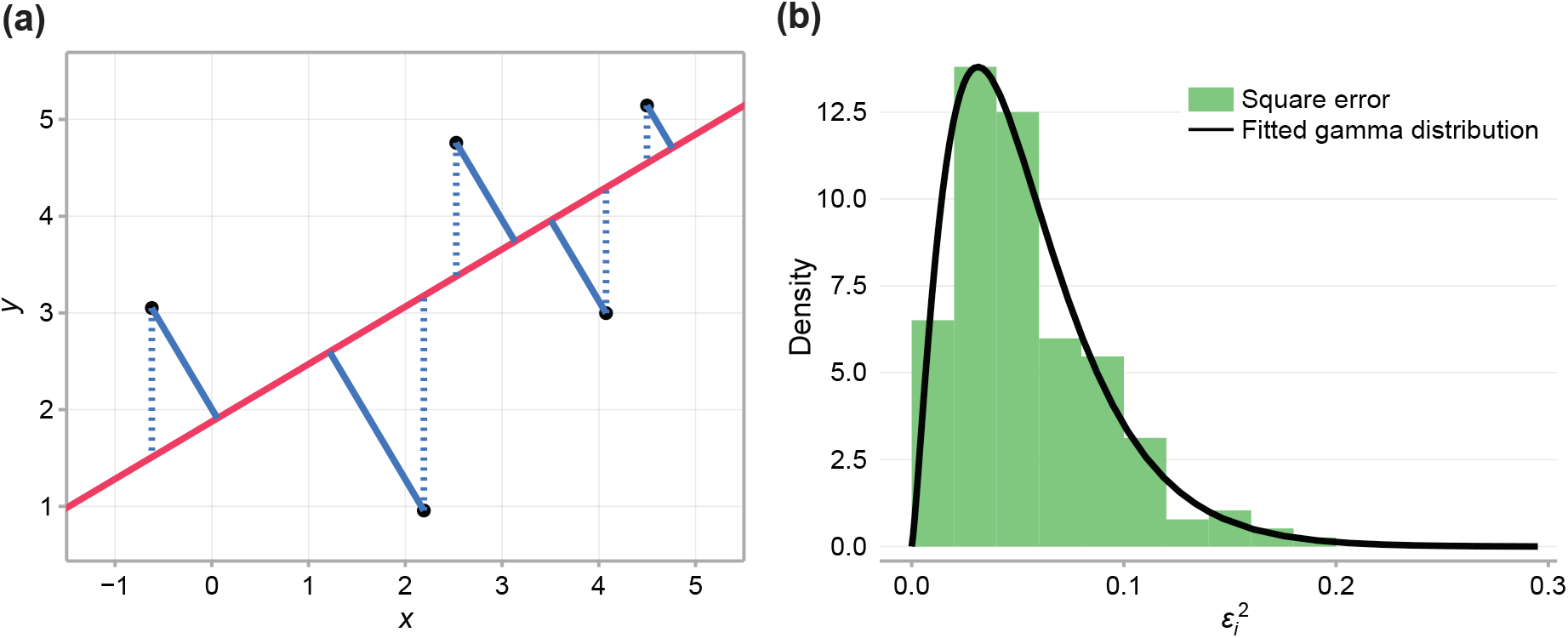
(a) Comparison between typical least-squares error (blue dashed), and total-least-squares error (blue solid). (b) Square error observed in the data and fitted gamma distribution

## Appendix 3

### Results for WM793b

**Figure A3.**
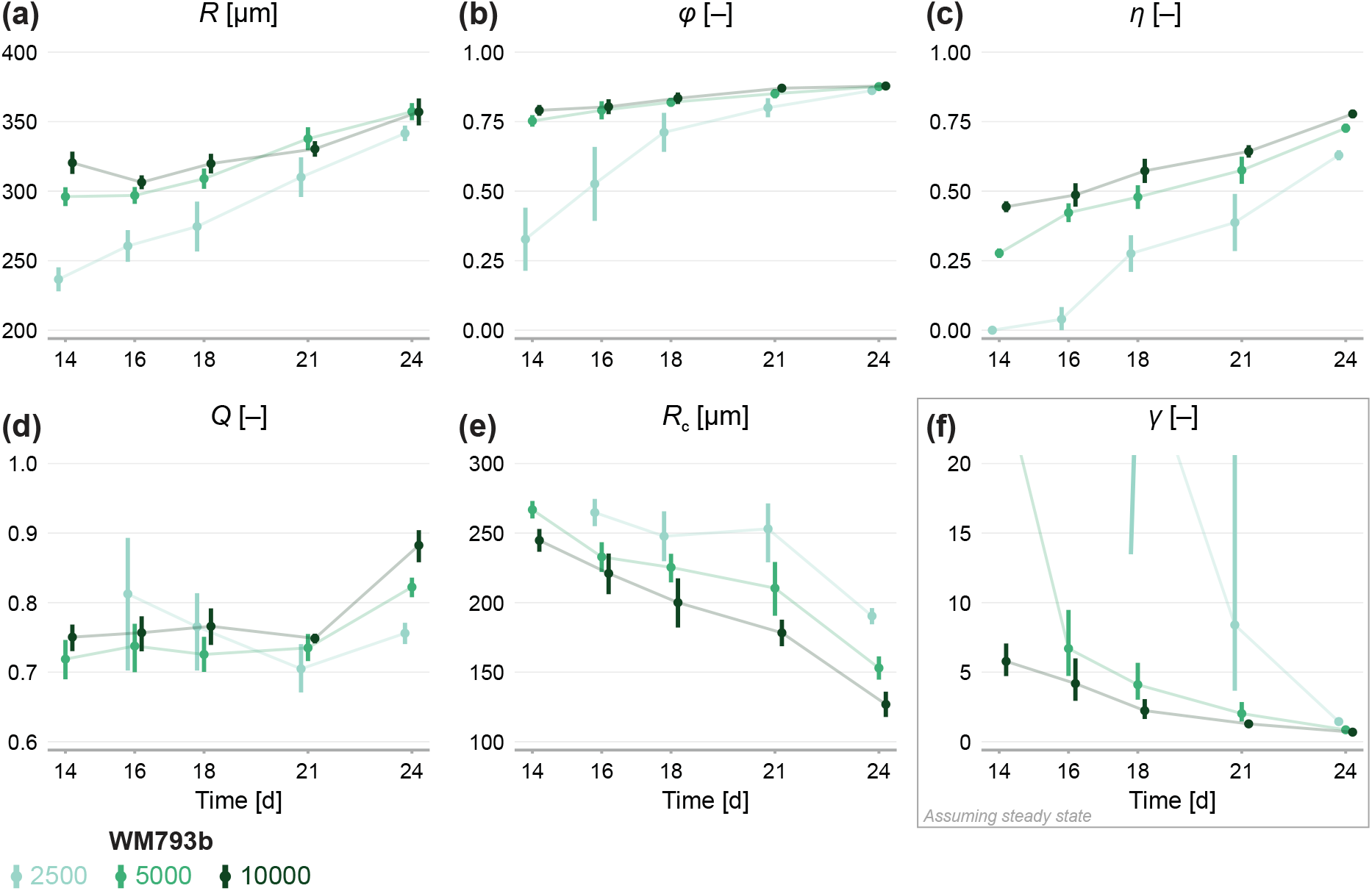
Estimates of parameters using the structural model with data from various time points. In (a–c), parameters are the mean of each observation: (*R, ϕ, η*). In (d–e), parameters are those in the structural model: (*R, Q, R*_c_). In (f), estimates of *γ* are obtained by calibrating observations to the steady-state model. As estimates *Q* and *R*_c_ can be derived from the structural model, which applies at any time during phase 3, we expect to see consistent estimates across observation times. Given that WM793b spheroids initiated with 2500 cells do not reach phase 3 until day 14, we exclude day 12 for these spheroids from the mathematical analysis. As estimates of *γ* can only be derived from the steady-state model, which assumes the outer radius is no longer increasing, we only expect consistency for later observation days. Bars indicate an approximate 95% confidence interval.

## Footnotes

1 Implemented to within machine precision using `Roots.jl`

2 Implemented by finding the eigenvalues of the characteristic matrix using `Polynomials.jl`

## References

[1] Hirschhaeuser F, Menne H, Dittfeld C, West J, Mueller-Klieser W, Kunz-Schughart LA. 2010. Multicellular tumor spheroids: An underestimated tool is catching up again. Journal of Biotechnology 148:3–15. doi:10.1016/j.jbiotec.2010.01.012.

[2] Cui X, Hartanto Y, Zhang H. 2017. Advances in multicellular spheroids formation. Journal of The Royal Society Interface 14:20160877. doi:10.1098/rsif.2016.0877.

[3] Smalley K, Lioni M, Noma K, Haass NK, Herlyn M. 2007. In vitro three-dimensional tumor microenvironment models for anticancer drug discovery. Expert Opinion on Drug Discovery 3:1–10. doi:10.1517/17460441.3.1.1.

[4] Santiago-Walker A, Li L, Haass N, Herlyn M. 2009. Melanocytes: from morphology to application. Skin Pharmacology and Physiology 22:114–121. doi:10.1159/000178870.

[5] Alexander S, Friedl P. 2012. Cancer invasion and resistance: interconnected processes of disease progression and therapy failure. Trends in Molecular Medicine 18:13–26. doi:10.1016/j.molmed.2011.11.003.

[6] LaBarbera DV, Reid BG, Yoo BH. 2012. The multicellular tumor spheroid model for high-throughput cancer drug discovery. Expert Opinion on Drug Discovery 7:819–830. doi:10.1517/17460441.2012.708334.

[7] Loessner D, Flegg JA, Byrne HM, Clements JA, Hutmacher DW. 2013. Growth of confined cancer spheroids: a combined experimental and mathematical modelling approach. Integrative Biology 5:597–605. doi:10.1039/c3ib20252f.

[8] Beaumont KA, Anfosso A, Ahmed F, Weninger W, Haass NK. 2015. Imaging-and flow cytometry-based analysis of cell position and the cell cycle in 3D melanoma spheroids. Journal of Visualized Experiments 106:e53486. doi:10.3791/53486.

[9] Langhans SA. 2018. Three-dimensional in vitro cell culture models in drug discovery and drug repositioning. Frontiers in Pharmacology 9:6. doi:10.3389/fphar.2018.00006.

[10] Theard PL, Sheffels E, Sealover NE, Linke AJ, Pratico DJ, Kortum RL. 2020. Marked synergy by vertical inhibition of EGFR signaling in nsclc spheroids shows sos1 is a therapeutic target in EGFR-mutated cancer. eLife 9:e58204. doi:10.7554/eLife.58204.

[11] Ivascu A, Kubbies M. 2006. Rapid generation of single-tumor spheroids for high-throughput cell function and toxicity analysis. Journal of Biomolecular Screening 11:922–932. doi:10.1177/1087057106292763.

[12] Friedrich J, Seidel C, Ebner R, Kunz-Schughart LA. 2009. Spheroid-based drug screen: considerations and practical approach. Nature Protocols 4:309–324. doi:10.1038/nprot.2008.226.

[13] Eilenberger C, Rothbauer M, Selinger F, Gerhartl A, Jordan C, Harasek M, Schdl B, Grillari J, Weghuber J, Neuhaus W, Kpc S, Ertl P. 2021. A microfluidic multisize spheroid array for multiparametric screening of anticancer drugs and bloodbrain barrier transport properties. Advanced Science 8:2004856. doi:10.1002/advs.202004856.

[14] Mark C, Grundy TJ, Strissel PL, Bhringer D, Grummel N, Gerum R, Steinwachs J, Hack CC, Beckmann MW, Eckstein M, Strick R, O’Neill GM, Fabry B. 2020. Collective forces of tumor spheroids in three-dimensional biopolymer networks. eLife 9:e51912. doi:10.7554/elife.51912.

[15] Folkman J, Hochberg M. 1973. Self-regulation of growth in three dimensions. The Journal of Experimental Medicine 138:745–753. doi:10.1084/jem.138.4.745.

[16] Greenspan HP. 1972. Models for the growth of a solid tumor by diffusion. Studies in Applied Mathematics 51:317–340. doi:10.1002/sapm1972514317.

[17] Adam J, Maggelakis S. 1990. Diffusion regulated growth characteristics of a spherical prevascular carcinoma. Bulletin of Mathematical Biology 52:549–582. doi:10.1016/s0092-8240(05)80362-3.

[18] Groebe K, Mueller-Klieser W. 1996. On the relation between size of necrosis and diameter of tumor spheroids. International Journal of Radiation Oncology, Biology, Physics 34:395–401. doi:10.1016/0360-3016(95)02065-9.

[19] Byrne HM, Chaplain MAJ. 1998. Necrosis and apoptosis: distinct cell loss mechanisms in a mathematical model of avascular tumour growth. Journ1al of Theoretical Medicine 1:223–235. doi:10.1080/10273669808833021.

[20] Ward JP, King JR. 1999. Mathematical modelling of avascular-tumour growth. II: Modelling growth saturation. IMA Journal of Mathematics Applied in Medicine and Biology 16:171–211. doi:10.1093/imammb/16.2.171.

[21] Araujo RP, McElwain DLS. 2004. A history of the study of solid tumour growth: The contribution of mathematical modelling. Bulletin of Mathematical Biology 66:1039–1091. doi:10.1016/j.bulm.2003.11.002.

[22] Wallace DI, Guo X. 2013. Properties of tumor spheroid growth exhibited by simple mathematical models. Frontiers in Oncology 3:51. doi:10.3389/fonc.2013.00051.

[23] Sarapata EA, de Pillis LG. 2014. A comparison and catalog of intrinsic tumor growth models. Bulletin of Mathematical Biology 76:2010 2024. doi:10.1007/s11538-014-9986-y.

[24] Flegg JA, Nataraj N. 2019. Mathematical modelling and avascular tumour growth. Resonance 24:313–325. doi:10.1007/s12045-019-0782-8.

[25] Murphy RJ, Browning AP, Gunasingh G, Haass NK, Simpson MJ. 2021. Designing and interpreting 4D tumour spheroid experiments. bioRxiv page 2021.08.18.456910. doi:10.1101/2021.08.18.456910.

[26] Gomes A, Guillaume L, Grimes DR, Fehrenbach J, Lobjois V, Ducommun B. 2016. Oxygen partial pressure is a rate-limiting parameter for cell proliferation in 3D spheroids grown in physioxic culture condition. PLOS ONE 11:e0161239. doi:10.1371/journal.pone.0161239.

[27] Herlyn M, Thurin J, Balaban G, Bennicelli JL, Herlyn D, Elder DE, Bondi E, Guerry D, Nowell P, Clark WH. 1985. Characteristics of cultured human melanocytes isolated from different stages of tumor progression. Cancer Research 45:5670–5676.

[28] Herlyn M. 1990. Human melanoma: Development and progression. Cancer and Metastasis Reviews 9:101–112. doi:10.1007/bf00046337.

[29] Sakaue-Sawano A, Kurokawa H, Morimura T, Hanyu A, Hama H, Osawa H, Kashiwagi S, Fukami K, Miyata T, Miyoshi H, Imamura T, Ogawa M, Masai H, Miyawaki A. 2008. Visualizing spatiotemporal dynamics of multicellular cell-cycle progression. Cell 132:487–498. doi:10.1016/j.cell.2007.12.033.

[30] Haass NK, Beaumont KA, Hill DS, Anfosso A, Mrass P, Munoz MA, Kinjyo I, Weninger W. 2014. Real-time cell cycle imaging during melanoma growth, invasion, and drug response. Pigment Cell & Melanoma Research 27:764–776. doi:10.1111/pcmr.12274.

[31] Kienzle A, Kurch S, Schlder J, Berges C, Ose R, Schupp J, Tuettenberg A, Weiss H, Schultze J, Winzen S, Schinnerer M, Koynov K, Mezger M, Haass NK, Tremel W, Jonuleit H. 2017. Dendritic mesoporous silica nanoparticles for pHstimuliresponsive drug delivery of TNFAlpha. Advanced Healthcare Materials 6:1700012. doi:10.1002/adhm.201700012.

[32] Spoerri L, Gunasingh G, Haass NK. 2021. Fluorescence-based quantitative and spatial analysis of tumour spheroids: a proposed tool to predict patient-specific therapy response. Frontiers in Digital Health 3:668390. doi:10.3389/fdgth.2021.668390.

[33] Spoerri L, Beaumont KA, Anfosso A, Haass NK. 2017. Real-time cell cycle imaging in a 3D cell culture model of melanoma. Methods in Molecular Biology 1612:401–416. doi:10.1007/978-1-4939-7021-629.

[34] Weiswald LB, Bellet D, Dangles-Marie V. 2015. Spherical cancer models in tumor biology. Neoplasia 17:1–15. doi:10.1016/j.neo.2014.12.004.

[35] Masuda F, Ishii M, Mori A, Uehara L, Yanagida M, Takeda K, Saitoh S. 2016. Glucose restriction induces transient G2 cell cycle arrest extending cellular chronological lifespan. Scientific Reports 6:19629. doi:10.1038/srep19629.

[36] Lehmann EL, Fienberg S, Casella G. 1998. Theory of Point Estimation, Second Edition. Springer, Secaucus. doi:10.1007/b98854.

[37] Raue A, Kreutz C, Maiwald T, Bachmann J, Schilling M, Klingmller U, Timmer J. 2009. Structural and practical identifiability analysis of partially observed dynamical models by exploiting the profile likelihood. Bioinformatics 25:1923–1929. doi:10.1093/bioinformatics/btp358.

[38] Pawitan Y. 2013. In all likelihood: statistical modelling and inference using likelihood. Oxford University Press, Oxford. ISBN 9780199671229.

[39] Browning AP, Maclaren OJ, Buenzli PR, Lanaro M, Allenby MC, Woodruff MA, Simpson MJ. 2021. Model-based data analysis of tissue growth in thin 3D printed scaffolds. Journal of Theoretical Biology doi:10.1016/j.jtbi.2021.110852.

[40] Ward JP, King JR. 1997. Mathematical modelling of avascular-tumour growth. IMA Journal of Mathematics Applied in Medicine and Biology 14:39–69. PMID: 9080687.

[41] Roose T, Chapman SJ, Maini PK. 2007. Mathematical models of avascular tumour growth. SIAM Review 49:179–208. doi:10.1137/S0036144504446291.

[42] Byrne HM. 2010. Dissecting cancer through mathematics: from the cell to the animal model. Nature Reviews Cancer 10:221–230. doi:10.1038/nrc2808.

[43] Bull JA, Mech F, Quaiser T, Waters SL, Byrne HM. 2020. Mathematical modelling reveals cellular dynamics within tumour spheroids. PLOS Computational Biology 16:e1007961. doi:10.1371/journal.pcbi.1007961.

[44] Gutenkunst RN, Waterfall JJ, Casey FP, Brown KS, Myers CR, Sethna JP. 2007. Universally sloppy parameter sensitivities in systems biology models. PLOS Computational Biology 3:e189. doi:10.1371/journal.pcbi.0030189.

[45] Gbor A, Banga JR. 2015. Robust and efficient parameter estimation in dynamic models of biological systems. BMC Systems Biology 9:74. doi:10.1186/s12918-015-0219-2.

[46] Raman DV, Anderson J, Papachristodoulou A. 2017. Delineating parameter unidentifiabilities in complex models. Physical Review E 95:032314. doi:10.1103/physreve.95.032314.

[47] Hoek KS, Schlegel NC, Brafford P, Sucker A, Ugurel S, Kumar R, Weber BL, Nathanson KL, Phillips DJ, Herlyn M, Schadendorf D, Dummer R. 2006. Metastatic potential of melanomas defined by specific gene expression profiles with no BRAF signature. Pigment Cell Research 19:290–302. doi:10.1111/j.1600-0749.2006.00322.x.

[48] Smalley KS, Contractor R, Haass NK, Kulp AN, Atilla-Gokcumen GE, Williams DS, Bregman H, Flaherty KT, Soengas MS, Meggers E, Herlyn M. 2007. An organometallic protein kinase inhibitor pharmacologically activates p53 and induces apoptosis in human melanoma cells. Cancer Research 67:209–217. doi:10.1158/0008-5472.can-06-1538.

[49] Smalley KSM, Contractor R, Haass NK, Lee JT, Nathanson KL, Medina CA, Flaherty KT, Herlyn M. 2007. Ki67 expression levels are a better marker of reduced melanoma growth following MEK inhibitor treatment than phospho-ERK levels. British Journal of Cancer 96:445–449. doi:10.1038/sj.bjc.6603596.

[50] Browning AP, Murphy RJ. 2021. Image processing algorithm to identify structure of tumour spheroids with cell cycle labelling. Zenodo doi:10.5281/zenodo.5121093.

[51] Mathworks. Texture analysis. https://au.mathworks.com/help/images/texture-analysis-1.html. Accessed: August 4, 2021.

[52] Laurent J, Frongia C, Cazales M, Mondesert O, Ducommun B, Lobjois V. 2013. Multicellular tumor spheroid models to explore cell cycle checkpoints in 3D. BMC Cancer 13:73. doi:10.1186/1471-2407-13-73.

[53] Murphy KC, Hung BP, Browne-Bourne S, Zhou D, Yeung J, Genetos DC, Leach JK. 2017. Measurement of oxygen tension within mesenchymal stem cell spheroids. Journal of The Royal Society Interface 14:20160851. doi:10.1098/rsif.2016.0851.

[54] Browning AP, Haridas P, Simpson MJ. 2019. A Bayesian sequential learning framework to parameterise continuum models of melanoma invasion into human skin. Bulletin of Mathematical Biology 81:676—698. doi:10.1007/s11538-018-0532-1.

[55] Hines KE, Middendorf TR, Aldrich RW. 2014. Determination of parameter identifiability in nonlinear biophysical models: A Bayesian approach. The Journal of General Physiology 143:401–16. doi:10.1085/jgp.201311116.

[56] Simpson MJ, Baker RE, Vittadello ST, Maclaren OJ. 2020. Practical parameter identifiability for spatio-temporal models of cell invasion. Journal of The Royal Society Interface 17:20200055. doi:10.1098/rsif.2020.0055.

[57] Powell MJD. 2009. The BOBYQA algorithm for bound constrained optimization without derivatives. Technical report. Department of Applied Mathematics and Theoretical Physics. Cambridge, England.

[58] Johnson SG. 2021. The NLopt module for Julia. https://github.com/JuliaOpt/NLopt.jl.

[59] Boiger R, Hasenauer J, Hro S, Kaltenbacher B. 2016. Integration based profile likelihood calculation for PDE constrained parameter estimation problems. Inverse Problems 32:125009. doi:10.1088/0266-5611/32/12/125009.

[60] Revels J, Lubin M, Papamarkou T. 2016. Forward-mode automatic differentiation in Julia. arXiv. https://arxiv.org/abs/1607.07892.

[61] Wang L, Cao J, Ramsay JO, Burger DM, Laporte CJL, Rockstroh JK. 2014. Estimating mixed-effects differential equation models. Statistics and Computing 24:111–121. doi:10.1007/s11222-012-9357-1.

[62] Hasenauer J, Hasenauer C, Hucho T, Theis FJ. 2014. ODE constrained mixture modelling: a method for unraveling subpopulation structures and dynamics. PLOS Computational Biology 10:e1003686. doi:10.1371/journal.pcbi.1003686.

[63] Jin W, Spoerri L, Haass NK, Simpson MJ. 2021. Mathematical model of tumour spheroid experiments with real-time cell cycle imaging. Bulletin of Mathematical Biology 83:44. doi:10.1007/s11538-021-00878-4.

[64] Lagergren JH, Nardini JT, Baker RE, Simpson MJ, Flores KB. 2020. Biologically-informed neural networks guide mechanistic modeling from sparse experimental data. PLOS Computational Biology 16:e1008462. doi:10.1371/journal.pcbi.1008462.

[65] Markovsky I, Huffel SV. 2007. Overview of total least-squares methods. Signal Processing 87:2283–2302. doi:10.1016/j.sigpro.2007.04.004.

[66] Welch BL. 1947. The generalization of ‘Student’s’ problem when several different population variances are involved. Biometrika 34:28. doi:10.2307/2332510.

