## Supplementary file 1 for "Quantitative analysis of tumour spheroid structure"

**Spheroid replicate count per condition.** We collect data on enough spheroids to ensure at least 10 for each initial seeding density and observation day (20 for comparing steady-state structure) are available. We then randomly subsample to ensure a consistent number of spheroids are analysed for each initial seeding density and observation day. The total number of spheroids, and the number in the subset, are given this table. Raw data (complete and the subset) are available on GitHub*.

| | Condition | | Day | | | | | | | | | | | Totals | |
|---|---|---|---|---|---|---|---|---|---|---|---|---|---|---|---|
| | | | 3 | 4 | 5 | 7 | 10 | 12 | 14 | 16 | 18 | 21 | 24 | | |
| All | 983b | 2500 | 6 | 12 | 12 | 20 | 16 | 16 | 13 | 17 | 13 | 26 | – | 151 | |
| | 983b | 5000 | 9 | 9 | 10 | 15 | 13 | 17 | 14 | 21 | 20 | 31 | – | 159 | 456 |
| | 983b | 10000 | 6 | 10 | 9 | 18 | 15 | 21 | 13 | 19 | 19 | 16 | – | 146 | |
| | 793b | 2500 | – | 5 | 10 | 22 | 28 | 27 | 19 | 18 | 20 | 15 | 23 | 187 | |
| | 793b | 5000 | – | 12 | 11 | 23 | 25 | 20 | 21 | 19 | 22 | 14 | 21 | 188 | 538 |
| | 793b | 10000 | – | 7 | 12 | 18 | 25 | 23 | 15 | 17 | 21 | 5 | 20 | 163 | |
| Subset | 983b | 2500 | 6 | 10 | 10 | 10 | 10 | 10 | 10 | 10 | 10 | 20 | – | 106 | |
| | 983b | 5000 | 9 | 9 | 10 | 10 | 10 | 10 | 10 | 10 | 20 | 10 | – | 108 | 318 |
| | 983b | 10000 | 6 | 10 | 9 | 10 | 10 | 10 | 10 | 10 | 19 | 10 | – | 104 | |
| | 793b | 2500 | – | 5 | 10 | 10 | 10 | 10 | 10 | 10 | 10 | 10 | 20 | 105 | |
| | 793b | 5000 | – | 10 | 10 | 10 | 10 | 10 | 10 | 10 | 10 | 10 | 20 | 110 | 317 |
| | 793b | 10000 | – | 7 | 10 | 10 | 10 | 10 | 10 | 10 | 10 | 5 | 20 | 102 | |

*github.com/ap-browning/Spheroids
