## Supplementary file 2 for "Quantitative analysis of tumour spheroid structure"

**Subset of spheroid cross-sectional confocal images.** We collect data on enough spheroids to ensure at least 10 for each initial seeding density and observation day (20 for comparing steady-state structure) are available. We then randomly subsample to ensure a consistent number of spheroids are analysed for each initial seeding density and observation day. In Fig. 1 to Fig. 6 we show a random subset of 10 spheroids from the complete data set for each condition, from days 7 to day 21 (WM983b) and day 24 (WM793b). Raw data (complete and the subset) are available on GitHub*.

#### WM983b (seeded with 2500 cells)

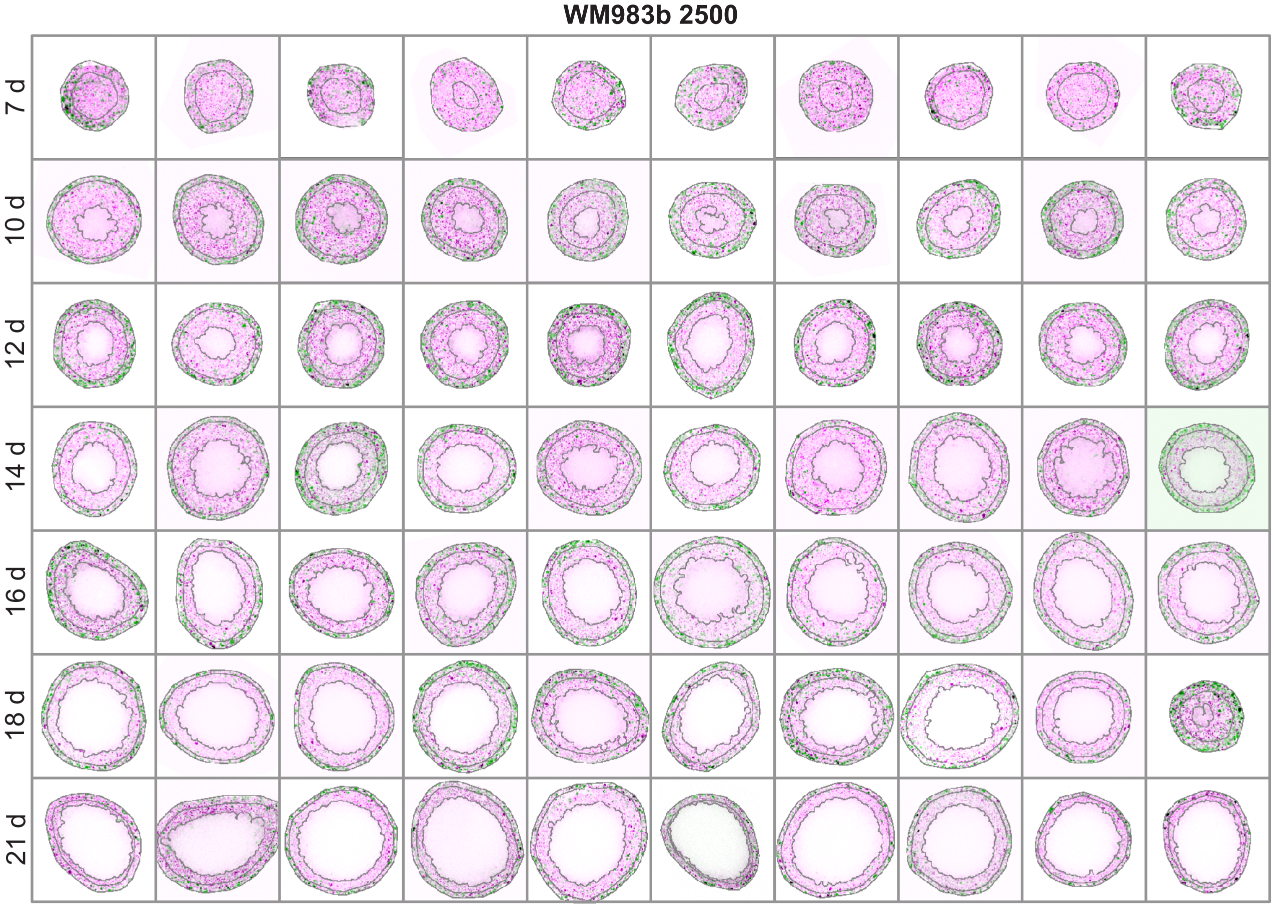


**Figure 1**

*github.com/ap-browning/Spheroids

### WM983b (seeded with 5000 cells)

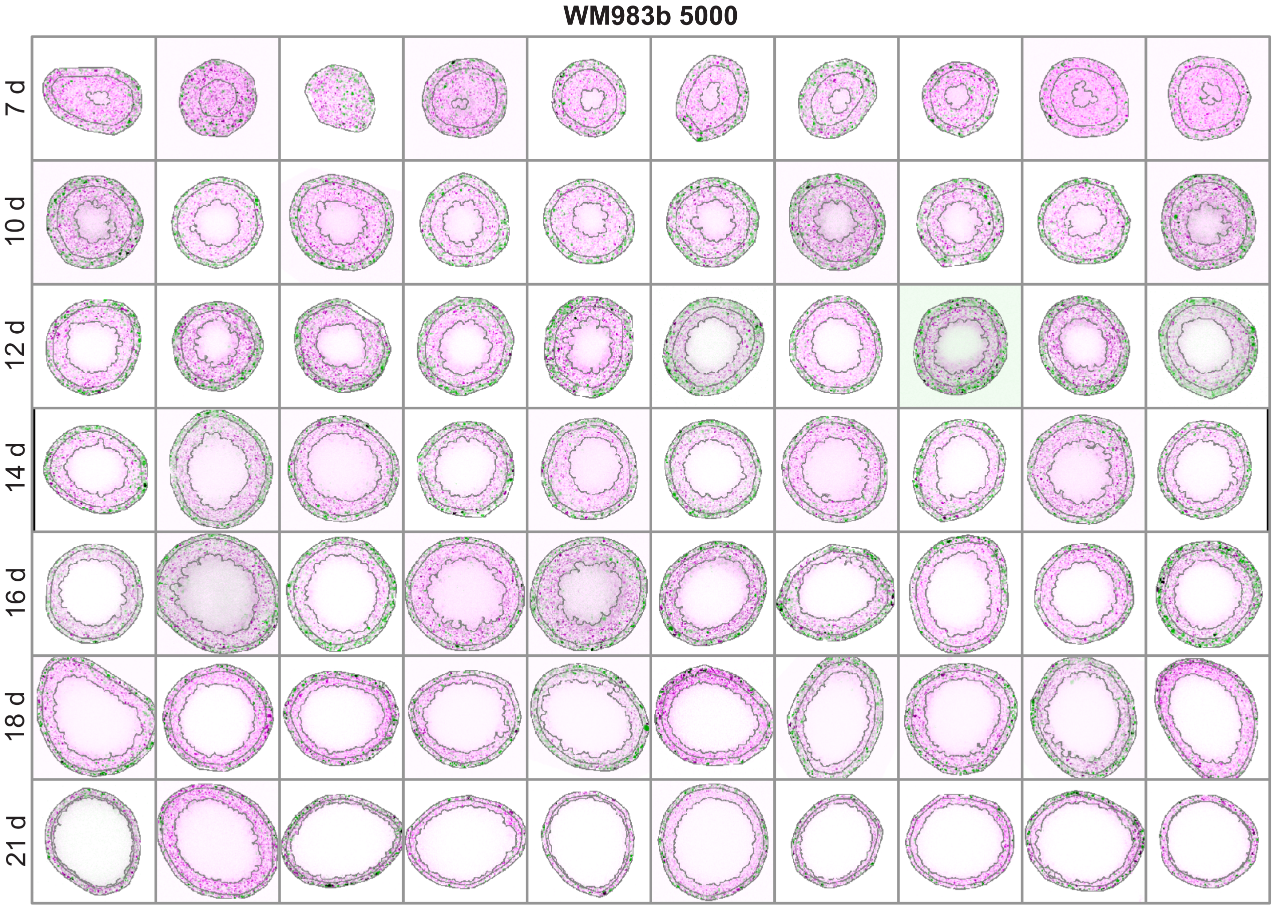


**Figure 2**

### WM983b (seeded with 10000 cells)

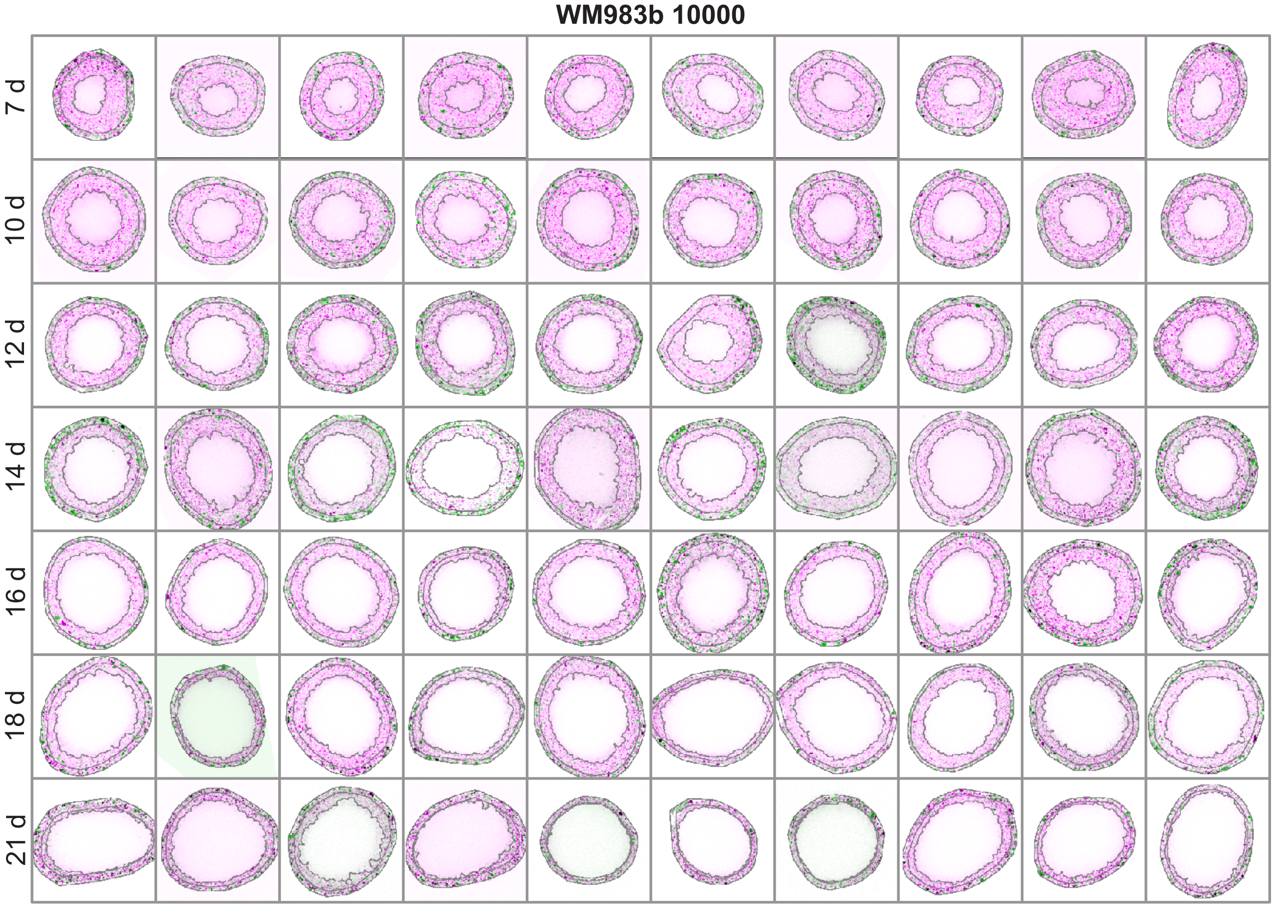


**Figure 3**

**WM793b (seeded with 2500 cells)**

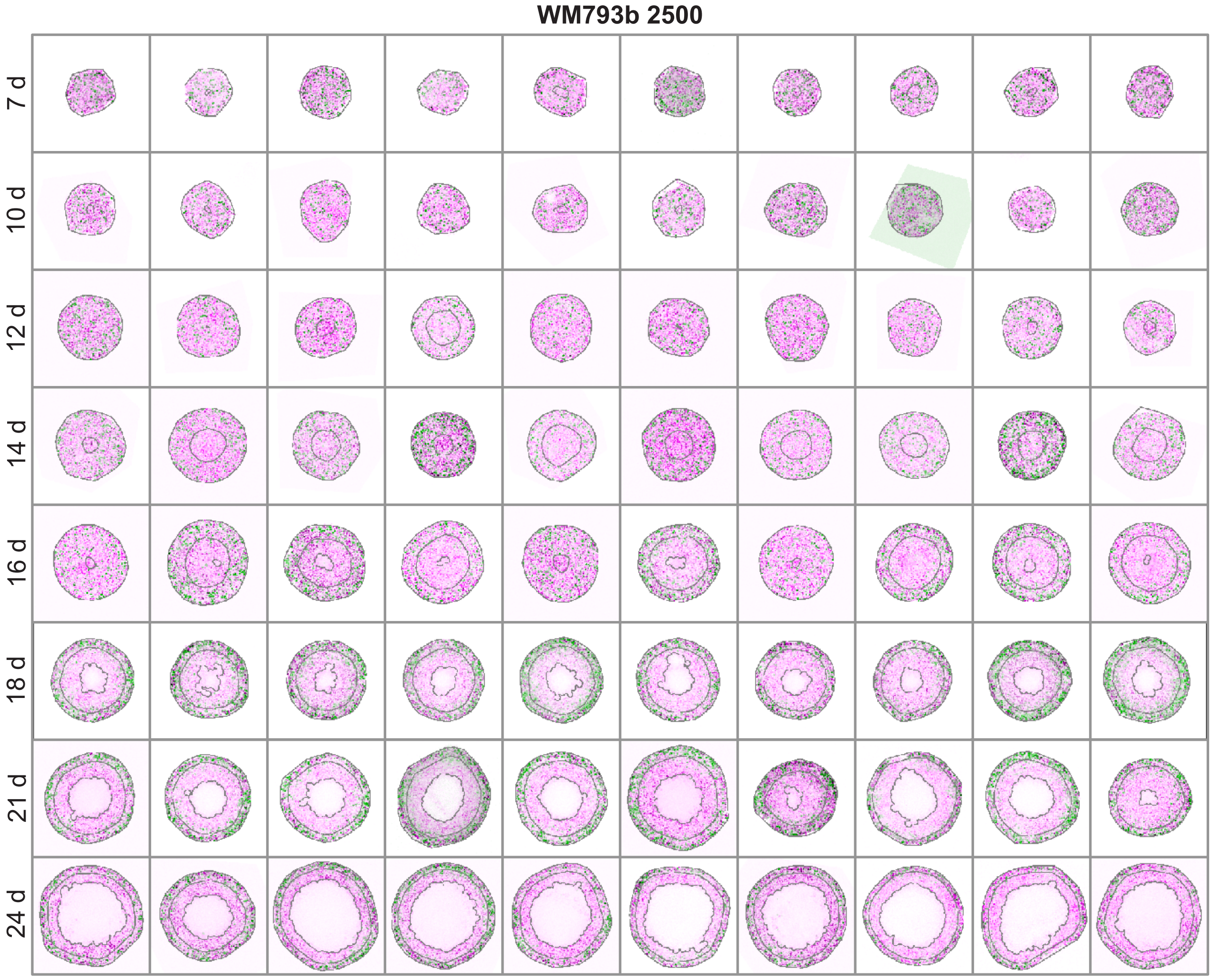


**Figure 4**

### WM793b (seeded with 5000 cells)

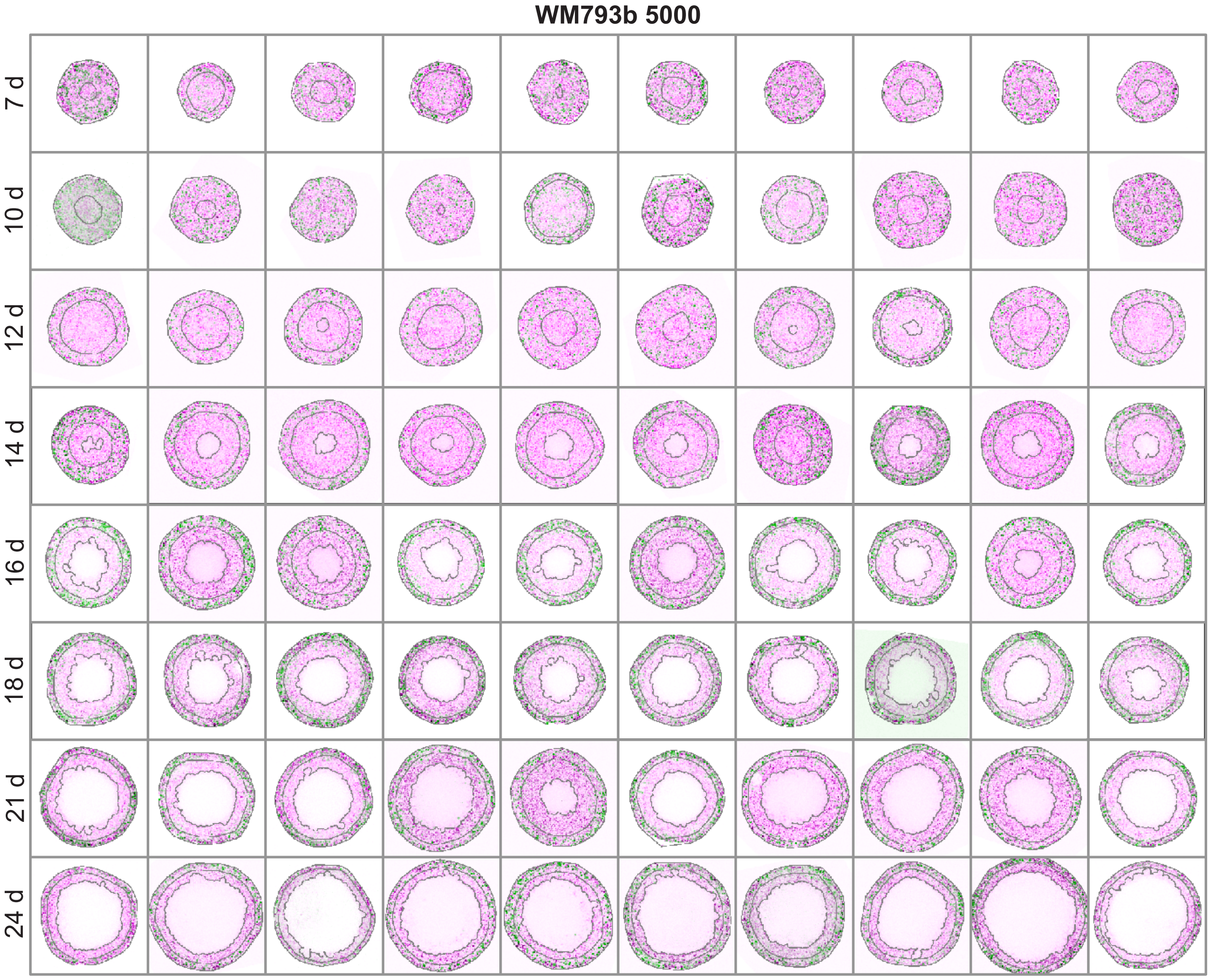


**Figure 5**

**WM793b (seeded with 10000 cells)**

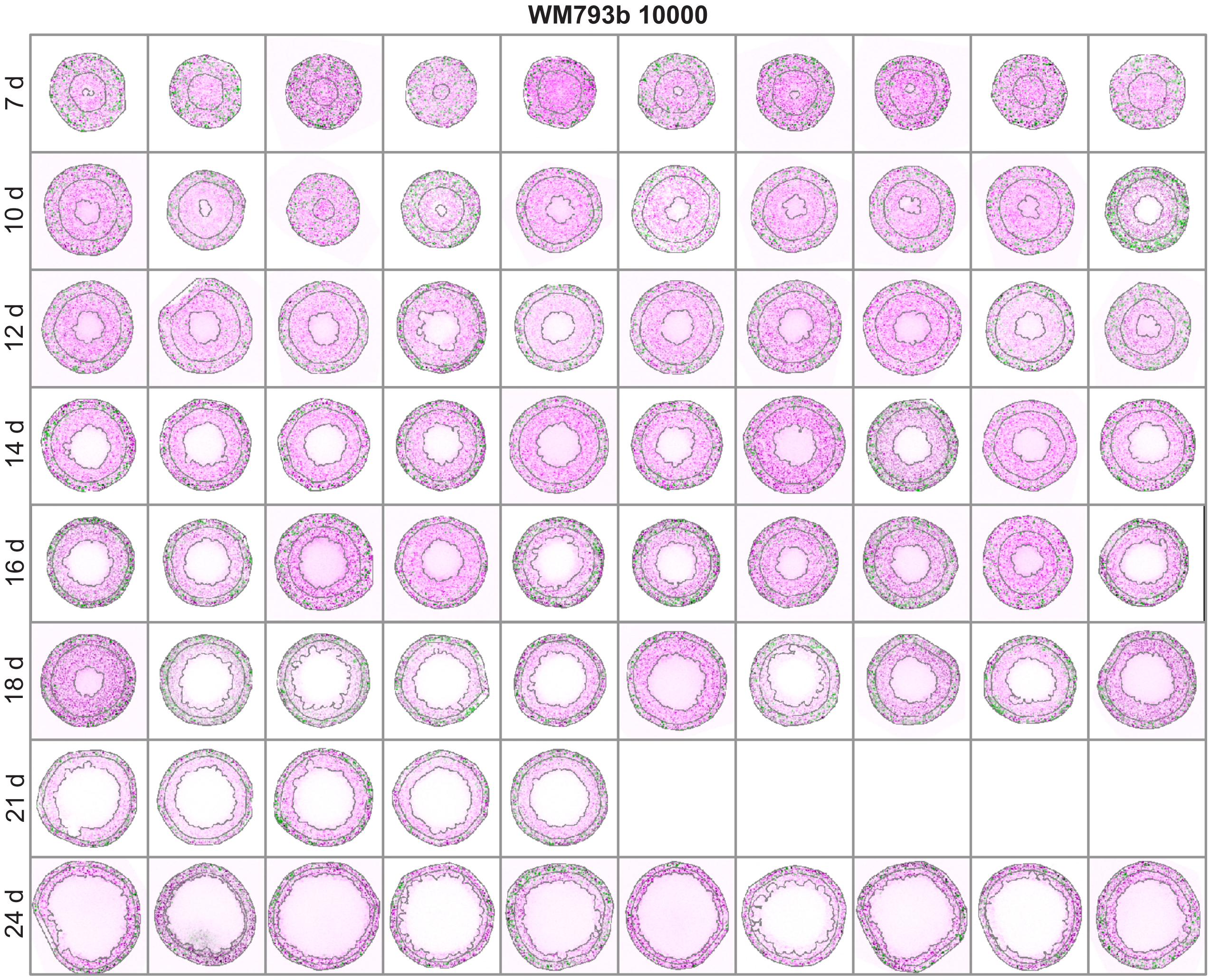


**Figure 6**
