## Supplementary file 3 for "Quantitative analysis of tumour spheroid structure"

**Steady-state analysis for WM983b cells using day 21 data.** In the main document, steady-state results in Figure 5 are produced by comparing data at day 21 (spheroids seeded with 2500 cells) to day 18 (spheroids seeded with 5000 and 10000 cells). Here, we reproduce the steady-state results in Figure 5 using data from day 21 from all initial seeding densities. However, we note that between days 18 and 21 spheroids seeded with 5000 and 10000 cells are observed to enter a fourth phase of decay, which is not captured by the mathematical model.

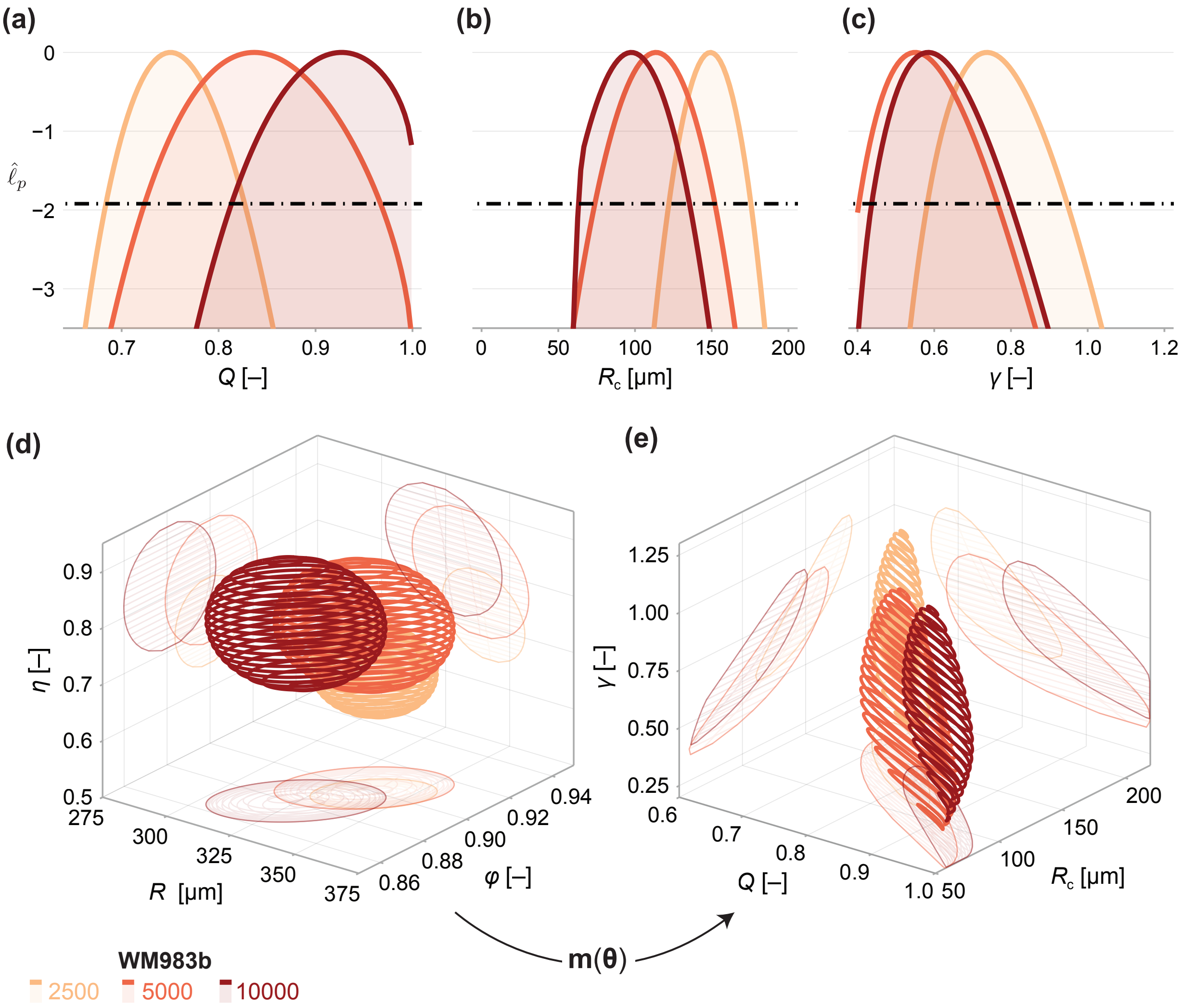


**Figure 1.** Comparison of WM983b spheroids between each initial seeding density at day 21. (a–c) Profile likelihoods for each parameter. (d) 95% confidence region for the full parameter space. 95% confidence regions for (d) the mean of each observation at steady state $(\bar{R}, \bar{\phi}, \bar{\eta})$ and (e) the model parameters $(Q, R_c, \gamma)$.
